# Stm1 regulates Ifh1 activity revealing crosstalk between ribosome biogenesis and ribosome dormancy

**DOI:** 10.1101/2024.04.22.590536

**Authors:** Eliana Bianco, Martina Bonassera, Federico Uliana, Janny Tilma, Martin Winkler, Sevil Zencir, Benjamin Albert, Michaela Oborská-Oplová, Reinhard Dechant, Jannik Hugener, Vikram Govind Panse, Martin Pilhofer, Philipp Kimmig, Matthias Peter

## Abstract

Ribosome abundance in changing environments is governed by biogenesis and degradation, but the underlying mechanisms regulating these opposing processes remain unknown. Here we show that Suppressor of Tom1 (Stm1), a dormancy factor protecting cytosolic ribosomes during starvation, has an unexpected function to promote ribosome biogenesis during exponential growth conditions. Indeed, Stm1 transiently localizes to the nucleolus and engages with pre-ribosomal particles. Stm1 upregulates transcription of ribosomal protein genes by directly binding the activation domain (AD) of the transcription factor Ifh1. These novel Stm1 functions confer rapamycin-sensitivity and are mediated by its C-terminal intrinsically disordered region (IDR), which is dispensable for ribosome hibernation. We conclude that Stm1 regulates ribosome homeostasis linking ribosome biogenesis and ribosome dormancy.

## Introduction

Ribosomes are conserved molecular factories that decode the genetic information into proteins, providing new components required for cell growth. Composed of ribosomal RNAs (rRNAs) and ribosomal proteins (RPs), the 80S ribosome comprises small 40S and the large 60S subunits. Ribosome biogenesis is a complex process through which rRNAs and RPs are synthesized and coordinately assembled into functional, cytoplasmic ribosomes (1). During ribosome biogenesis, the primary 35S rRNA is transcribed by RNA polymerase I (Pol I) in the nucleolus and processed into mature 18S, 25S and 5.8S rRNA. In the nucleus, 5S rRNA, tRNAs and other short RNAs are transcribed by RNA Pol III, while RP messenger RNA (mRNA), small nuclear RNAs (snRNA) and small nucleolar RNAs (snoRNA) are transcribed by RNA polymerase II (Pol II). mRNAs are translated into proteins by ribosomes in the cytoplasm and most RPs then travel back to the nucleolus or nucleus to be assembled with rRNAs into new ribosomes aided by ribosome assembly factors (RiBi) (2–5)(**Fig 1A**). Altogether, over 200 biogenesis factors and more than 75 snoRNAs are involved in ribosome biogenesis, comprising one of the most energy consuming cellular processes. Not surprisingly, errors in ribosome biogenesis are linked to defects in cell growth, ribosomopathies and cancer (6–8).

**Figure 1:**
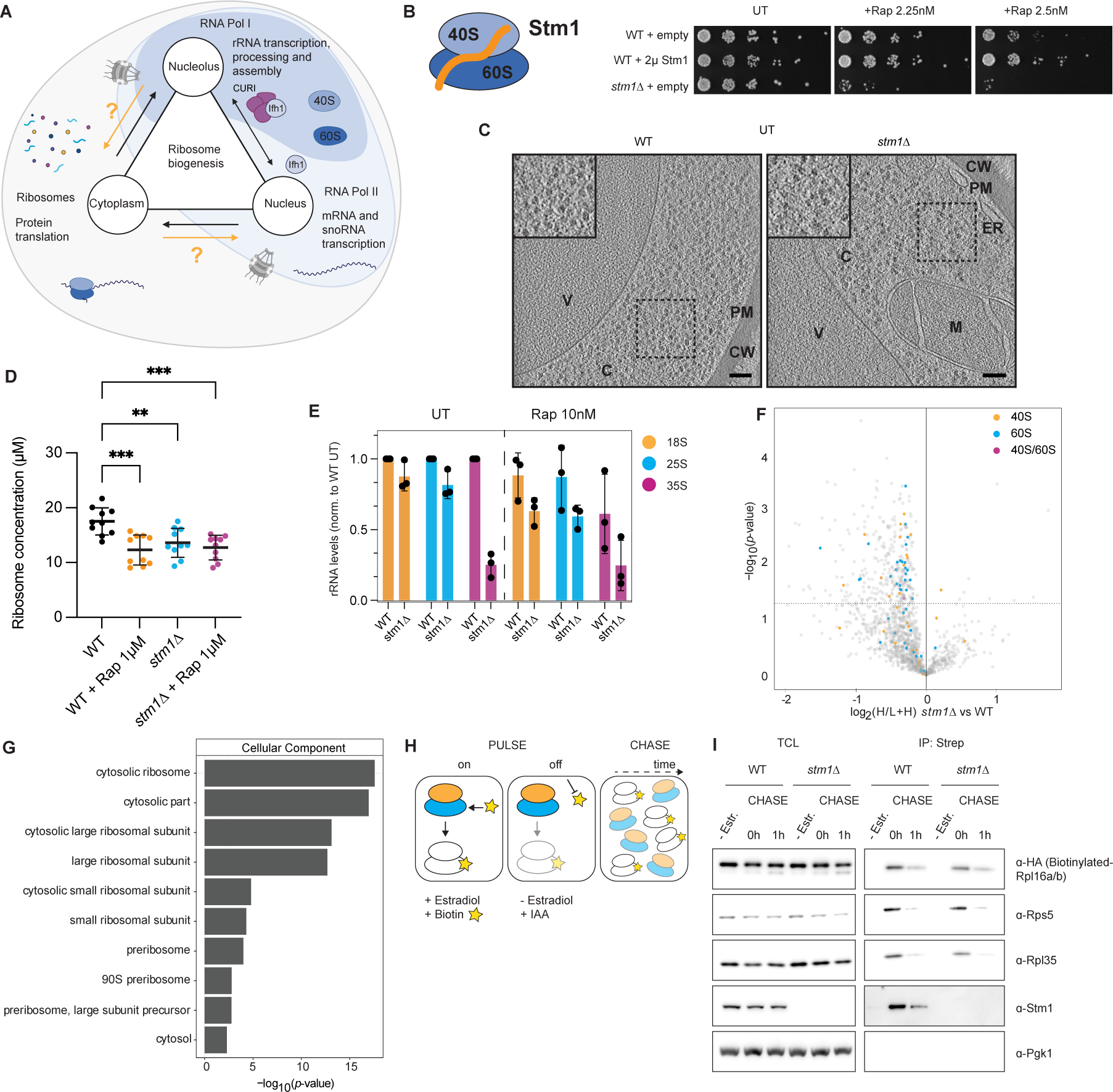
The ribosome hibernation factor Stm1 promotes ribosome biogenesis. **A)** Schematic representation how ribosome biogenesis is orchestrated across different cellular compartments, including nucleolus, nucleus and cytoplasm. rRNA transcription, processing and assembly with RiBi factors and RPs occurs in the nucleolus. In the nucleus, RNA Pol II transcribes mRNAs and snoRNAs, and RNA Pol III transcribes 5S rRNA, tRNAs and other non-coding RNAs. The CURI complex and the transcription factor Ifh1 are depicted between the nucleolus and nucleus, thereby fine-tuning the rate of RPG transcription. RPs are translated in the cytoplasm and then imported into the nucleolus or nucleus to be assembled into pre-RPs. Pre-RPs are exported to the cytoplasm, assemble as 80S mature ribosomes and engage in the translation cycle. Orange arrows and question marks indicate unknown crosstalk mechanisms linking cytoplasmic translation and ribosome biogenesis processes in nucleolus and nucleus. **B)** Schematic drawing (left panel) of Stm1 (orange) bound to a 80S ribosome composed of 40S (light blue) and 60S (dark blue) subunits. Serial dilution spotting assays (right panel) on synthetic auxotrophic SD-URA media to monitor growth of wild type (WT) and *stm1D* strains harboring an empty control plasmid (empty) or overexpressing Stm1 from a 2µ plasmid (2µ Stm1). Cells were either untreated (UT) or treated with rapamycin (Rap) at the indicated concentrations (2.25nM or 2.5nM) and incubated for 3 days. Representative image from N=3 independent experiments. **C)** Representative cryo-tomograms of untreated WT and *stm1Δ* cells, which were subjected to plunge freezing and focused ion beam (FIB)-milling prior to cryo-electron tomography. Shown are projections of 10.7nm-thick slices. Regions of interest (square dashed line) are shown magnified as an inset (solid black line). C: cytoplasm, ER: endoplasmic reticulum, M: mitochondrion, PM: plasma membrane V: vacuole. CW: cell wall. Scale bars: 100 nm. Additional cryo-tomograms are shown in **Sup 1E.** **D)** Quantification of ribosome concentration (µM) from cryo-tomograms of WT and *stm1Δ* cells, treated or not with rapamycin (Rap 1µM). Each dot represents the mean concentration of five independent measurements of ribosome concentration within one tomogram (workflow shown in **Sup 1D**). Data values are shown as mean with standard deviation. N=10 tomograms per experimental condition. The indicated *P*-values were calculated by one-way ANOVA test for multiple comparison analysis (Holm-Sidak). Significance: * p-value < 0.05; ** p-value < 0.01; *** p-value < 0.001. Note that there is no significant difference of the ribosome concentration observed in WT + Rap, *stm1Δ* and *stm1Δ* + Rap. **E)** Bar plot quantifying 18S (orange), 25S (blue), and 35S (purple) rRNA levels in WT and *stm1Δ* cells, treated (Rap) or not (UT) (left) with 10 nM rapamycin. Black dots represent independent quantifications of rRNA normalized to untreated WT controls. Mean with standard deviation are shown. Representative image of a rRNA gel (N=3 independent replicates) is shown in **Sup 1F**. **F)** Volcano plot of pulsed SILAC MS measurements comparing newly synthetized proteins in WT and *stm1Δ* cells. Protein levels are expressed as the ratio of proteins incorporating heavy lysine over total levels (log_2_ (H/L+H)). The significance is calculated from unpaired two-sided Student’s t-test. 40S subunit ribosomal proteins are labelled in orange, 60S proteins in blue and proteins shared by both subunits (40S/60S) highlighted in purple. N=4 independent replicates. **G)** Gene ontology enrichment calculated for downregulated proteins (*p-*value< 0.05) in *stm1Δ* cells. The significance of the enrichment is reported as the -log_10_ of *p*-value. **H)** *In vivo* pulse-chase proximity biotin ribosome labelling system. During the pulse phase (on), the 60S subunit (blue) protein Rpl16a/b-HA-TEV-AVI is biotinylated (yellow star) by the estradiol-inducible biotin ligase BirA. BirA degradation by the auxin inducible degron abolishes further biotin labelling (off). Biotinylated 60S ribosomes can be chased over time and in different experimental conditions and detected by streptavidin. This method allows to distinguish old (biotin-labelled) from newly synthesized ribosomes (no label) and thus to study ribosome turnover. **I)** Western blot of total cell lysates (TCL) and co-precipitated streptavidin-bound proteins (IP: Strep) from WT and *stm1Δ* cells expressing endogenously tagged Rpl16a/b-HA-TEV-AVI. Samples were left uninduced (-estradiol) or supplemented with 5nM estradiol for 2h to induce BirA expression. After washing out estradiol, samples were supplemented with biotin to induce biotin labelling of Rpl16a/b-HA-TEV-AVI and 1 mM auxin (IAA) to initiate BirA degradation. Samples were harvested right after the BirA biotinylation/degradation step (CHASE, 0h) or 1 h after BirA shut off and degradation (CHASE, 1h). Blots were probed with HA-antibodies (biotinylated Rpl16a/b-HA-TEV-AVI undergone TEV elution from Strep-beads), endogenous Rps5 and Rpl35, Stm1, and Pgk1 as a loading control. N=3 independent replicates. Note that reduction of ribosomal subunits in *stm1Δ* cells is comparable to WT controls.

Coordination of ribosome biosynthesis across cellular compartments is fundamental to maintain ribosome abundance (**Fig 1A**). Indeed, perturbing ribosome biogenesis causes proteotoxic stress and reduces cellular fitness (9, 10). How, then, are the various ribosome biosynthesis activities coordinated across the nucleolus, nucleus and cytoplasm to maintain ribosome homeostasis? Molecular dissection reveals signaling crosstalk between nucleolar rRNA transcription and maturation, and nuclear ribosomal protein gene (RPG) transcription involves the inhibitory function of the nucleolar CURI complex (11, 12) (**Fig 1A**). When RNA Pol I activity is downregulated, the UtpC subunits Utp22 and Rrp7, and the CK2 kinase subunit Ckb2 sequester the RPG transcription activator Interaction with Fork Head 1 (Ifh1) in the nucleolus, assembling into the CURI complex (11). Therefore, Ifh1 promoter binding activity is inhibited, and RPG transcription is downregulated (12). In contrast, evidence is lacking for cellular components and mechanisms that coordinate the cytoplasmic translation rate of RPs and RiBi with nucleolar Ifh1 activity and pre-RP assembly. Such a crosstalk mechanism is needed for cells to ensure ribosome homeostasis and link ribosome production to cytoplasmic translation in response to changing growth conditions. Conceptually, such cytoplasmic-nuclear-nucleolus crosstalk would require regulatory factors that exhibit nuclear and cytosolic localization, bind cytosolic ribosomes, and associate with pre-ribosomal particles and/or RPG transcription factors in the nucleus/nucleolus (**Fig 1A, question marks**).

The primary stimulus for ribosome biogenesis is nutrient availability, due to the central function of ribosomes in cell growth. In nutrient rich conditions the ribosome biogenesis pathways are active. In response to starvation the demand for new protein synthesis and ribosome production is reduced, and ribosome biogenesis in the nucleolus, nucleus and cytoplasm encompassing transcription, translation and rRNA maturation are rapidly downregulated and eventually switched off. The master molecular regulators governing ribosome biogenesis in response to nutrient availability are protein kinase A (PKA) and the target of rapamycin complex 1 (TORC1). These nutrient-responsive kinases promote transcription of RiBi factors and RPs by regulating the activity of transcription repressors and activators (13–15). TORC1 and PKA also repress catabolic processes such as degradation of macromolecules or macro-autophagy (16, 17). Nitrogen starvation or treatment with the macrolide rapamycin are potent inhibitors of TORC1, thereby repressing RiBi and RPG transcription and protein translation (18), and inducing autophagy and ribophagy, which specifically targets ribosomes for degradation (17, 19–22).

When nutrients are replenished following starvation, cells restart growth and resume protein translation. Thus, although ribosomes are degraded during starvation, growth cannot restart without available and functional ribosomes engaging in the translation cycle. Current evidence suggests that growth restart requires a specialized pool of protected ribosomes that are maintained in an inactive and dormant state during starvation (23, 24). Indeed, after rapamycin treatment cells preserve about 50-60 % of their ribosomes compared to ribosome abundance during nutrient rich conditions (25, 26). Specific ribosome dormancy factors have been identified in bacteria, which are upregulated upon adaptation to starvation and stress in bacteria (27, 28). Likewise, factors protecting eukaryotic ribosomes from degradation during starvation have been described, such as the functionally-conserved budding yeast Suppressor of Tom1 (Stm1) (24, 29, 30), its zebrafish and *Xenopus* homolog Habp4 (31) or its human homolog SERBP1 (32). Stm1 occupies the mRNA tunnel and contacts both 40S and 60S subunits of the 80S ribosome (**Fig 1B, left panel**) (30). Upon replenishment of nutrients, TORC1 directly phosphorylates Stm1, which then releases from dormant ribosomes allowing translation restart and resuming cell growth (24). Deletion of Stm1 sensitizes cells to rapamycin and compromises cell growth restart after nitrogen starvation (23), which has been attributed to ribosome depletion. Altogether, despite our detailed knowledge of ribosome biogenesis and insights into ribosome degradation/protection mechanisms, evidence is lacking for functional coupling to maintain ribosome homeostasis. Moreover, factors that regulate the abundance of ribosomes have yet to be described.

Here we report an unexpected nucleolar and nuclear function of Stm1 during exponential growth, distinct from its cytosolic dormancy function during starvation. During growth, Stm1 transiently shuttle into the nucleolus and interacts with pre-ribosomal particles and components of the CURI complex. Through its C-terminal intrinsically disordered region (IDR), Stm1 directly binds Ifh1 and promotes transcription of RPGs. Our results thus demonstrate that Stm1 fulfills a dual function. In the absence of stress, Stm1 shuttles into the nucleolus/nucleus and activates RPG production and ribosomes biogenesis. Upon nutrient starvation, Stm1 functions as a cytoplasmic dormancy factor to inactivate and protect a subset of mature ribosomes. Thus, Stm1 regulates ribosome homeostasis by coupling ribosome biosynthesis and degradation in response to nutrient availability.

## Results

### Stm1 promotes ribosome biogenesis during exponential growth

To understand the cellular function of the dormancy factor Stm1, we first used spotting assays to assess rapamycin sensitivity of wild-type, *STM1* knock-out (*stm1Δ)* and Stm1 overexpressing cells (2µ Stm1) (**Fig 1B**) (29). Since TORC1 controls cell growth through upregulation of transcription and translation machineries (33) we also tested *stm1Δ* sensitivity to specific translation inhibitors. As expected, *stm1Δ* cells were sensitive to rapamycin, while Stm1 overexpression confers resistance to higher rapamycin concentrations (**Fig 1B**). The sensitivity of wild-type and *stm1Δ* cells to the translation inhibitors cycloheximide (CHX) and hygromycin B (HygB) was comparable, whereas Stm1 overexpression showed increased toxicity (**Sup 1A**) (34). We conclude that Stm1 is essential in cells with reduced TORC1-activity and the rapamycin sensitivity of *stm1Δ* cells cannot be explained just by translational defect.

To examine Stm1 expression, we next analyzed its mRNA and protein levels. *STM1* mRNA is present in exponentially growing cells, and its expression is downregulated upon TORC1 inhibition or heat shock, similar to RiBi factors and RPs (**Sup 1B**). Likewise, Stm1 protein levels are abundant in rapidly growing cells, but decreased upon starvation (**Sup 1C**). The observed Stm1 expression pattern was a surprise, since bacterial hibernation factors are known to be upregulated upon adaptation to starvation and stress (27, 28). Moreover, Stm1 expression in nutrient-rich conditions suggests that the function of Stm1 may not be restricted to ribosome protection during nutrient starvation.

Since TORC1 promotes RP and RiBi expression, we compared the concentration of 80S cytosolic ribosomes in wild-type and *stm1Δ* cells using *in situ* cryo-electron tomography (cryoET). To this end, we developed a pipeline to segment and count the number of ribosomes per volume (**Sup 1D**). Consistent with previous results (25), the concentration of ribosomes in wild-type cells was ∼18µM and dropped to ∼12µM upon rapamycin treatment. In *stm1Δ* cells, the cytoplasmic ribosome concentration was significantly lower (∼14µM) compared to wild-type controls (∼18µM), and further reduced upon rapamycin treatment (∼13µM) (**Fig 1C and D, Sup 1E**). Since it is experimentally challenging to measure the volume of cytoplasm, we corroborated these findings by two additional methods. We quantified the levels of different rRNA species in wild-type and *stm1Δ* cells exposed or not to rapamycin. Indeed, 18S and 25S rRNA levels decreased by 10-20% in unchallenged *stm1Δ* cells and were further reduced by 40% upon TORC1 inhibition (**Fig 1E, Sup 1F**). Strikingly, the levels of unprocessed RNA Pol I transcript 35S were markedly diminished by approximately 75% in untreated *stm1Δ* cells or cells exposed to low rapamycin concentrations (**Fig 1E, Sup 1F**). Together, these results suggest that Stm1 is required for ribosome homeostasis and promotes ribosome biosynthesis in unstressed cells, potentially at early biogenesis steps.

As a second approach and to examine the mechanism of this unexpected Stm1 function, we assessed the cause of decreased ribosome numbers in *stm1Δ* cells. We measured the levels of newly synthesized proteins in wild-type and *stm1Δ* cells using pulsed-SILAC-based quantitative mass spectrometry (MS) analysis (35). Interestingly, deletion of Stm1 significantly decreases the expression levels of newly-produced proteins with an predominant decrease in ribosomal proteins (**Fig 1F and G, Sup 1G, Supplementary table 1**). Since *stm1Δ* does not affect the cellular growth rate (**Sup 1H**), this finding is consistent with the reduction of ribosome concentration (**Fig 1C and D, Sup 1E**) and rRNA levels (**Fig 1E**). Among the most downregulated proteins, we found Rpb5, a common component of RNA Pol I, RNA Pol II and RNA Pol III (36), which could explain the drop in 35S abundance (**Supplementary table 1**). Consistent with growth rate measurements (**Sup 1H**), upon exposure to CHX or HygB (**Sup 1A**) wild-type and *stm1Δ* cells incorporate puromycin at similar rates (**Sup 1I**), and in both strains Sui2 (eIF2-alpha) and Rps6 phosphorylation is comparable (**Sup 1J**). These results suggest Stm1 absence does not affect protein translation. Taken together, we conclude that dormancy factor Stm1 has an unexpected function in ribosome homeostasis in exponentially growing cells.

To assess whether decreased ribosome numbers in *stm1Δ* cells are caused by increased ribosome turnover, we developed a pulse-chase biotinylation proximity labelling system to specifically follow the fate of endogenously AVI-tagged mature cytosolic ribosomes *in vivo* (**Fig 1H**). Briefly, HA-TEV-AVI-tagged Rpl16a/b (37) are biotinylated by cytosolic BirA biotin ligase. The levels of BirA ligase are controlled by estradiol-induced transcription (38) and auxin-based protein degradation (39), thereby allowing rapid biotinylation at the AVI-tag on ribosomal proteins Rpl16a/b in a spatial and temporally controlled manner (**Sup 1K**). Importantly, this system allows isolating biotinylated cytoplasmic ribosomes by affinity purification and thus directly comparing their turnover in wild-type and *stm1Δ* cells by immunoblotting and Coomassie stain readouts. Interestingly, biotinylated ribosomal subunits Rpl16a/b, as well as Rpl35 and Rps5, were reduced to a similar extent in both strains after 1 hour of chase (1h) in exponential growth, indicating that Stm1 does not control ribosome stability in nutrient-rich conditions (**Fig 1I, Sup 1L**). These results indicate that the reduced ribosome concentration in *stm1Δ* cells does not result from increased ribosome degradation, but rather reduced ribosome biogenesis.

### Stm1 interacts with ribosome assembly factors in the nucleolus

To elucidate the mechanism of this novel Stm1 function in ribosome biogenesis, we stably integrated a single copy of fully functional 3xMyc-tagged Stm1 in the genome of *stm1Δ* cells (40). Stm1-3xMyc was then purified using anti-Myc affinity-beads, and interacting proteins were identified by mass-spectrometry (MS). To filter contaminants, we included cells expressing 3xMyc-tagged GFP. The quality control for the AP-MS experiments is reported in **Sup 2A, B, C**. As expected, Stm1-3xMyc pulled down ribosomal subunits and ribosome-interacting proteins, consistent with a role for Stm1 as a cytoplasmic hibernation factor. Surprisingly, the gene ontology (GO) enrichment analysis for “Cellular Component” and “Molecular Function” categories showed that RiBi factors predominantly residing in the nucleolus and/or nucleus (“preribosome”, “preribosome, large subunit processor” and “ribosome biogenesis”) are co-purified with Stm1 (**Fig 2A, B, C**, **Sup 2D, Supplementary table 2**). Two-hybrid analysis corroborated that Stm1 binds several nucleolar RiBi-factors, including other SSU components such as Utp18 and Utp22, in addition to Noc4 (**Fig 2D**). To further verify that Stm1 physically associates with nucleolar pre-RPs, we purified Nop7-TAP and Noc4-TAP particles belonging to the early steps of 60S (pre-66S) and 40S (small subunit processome, SSU) biogenesis respectively, and probed these complexes for the presence of untagged, endogenous Stm1. We found Stm1 specifically interacted with both pre-RPs (**Fig 2E**). We noticed that two upshifted Stm1 bands co-purified with Noc4-TAP but not Nop7-TAP particles, which may represent post-translational modifications.

**Figure 2:**
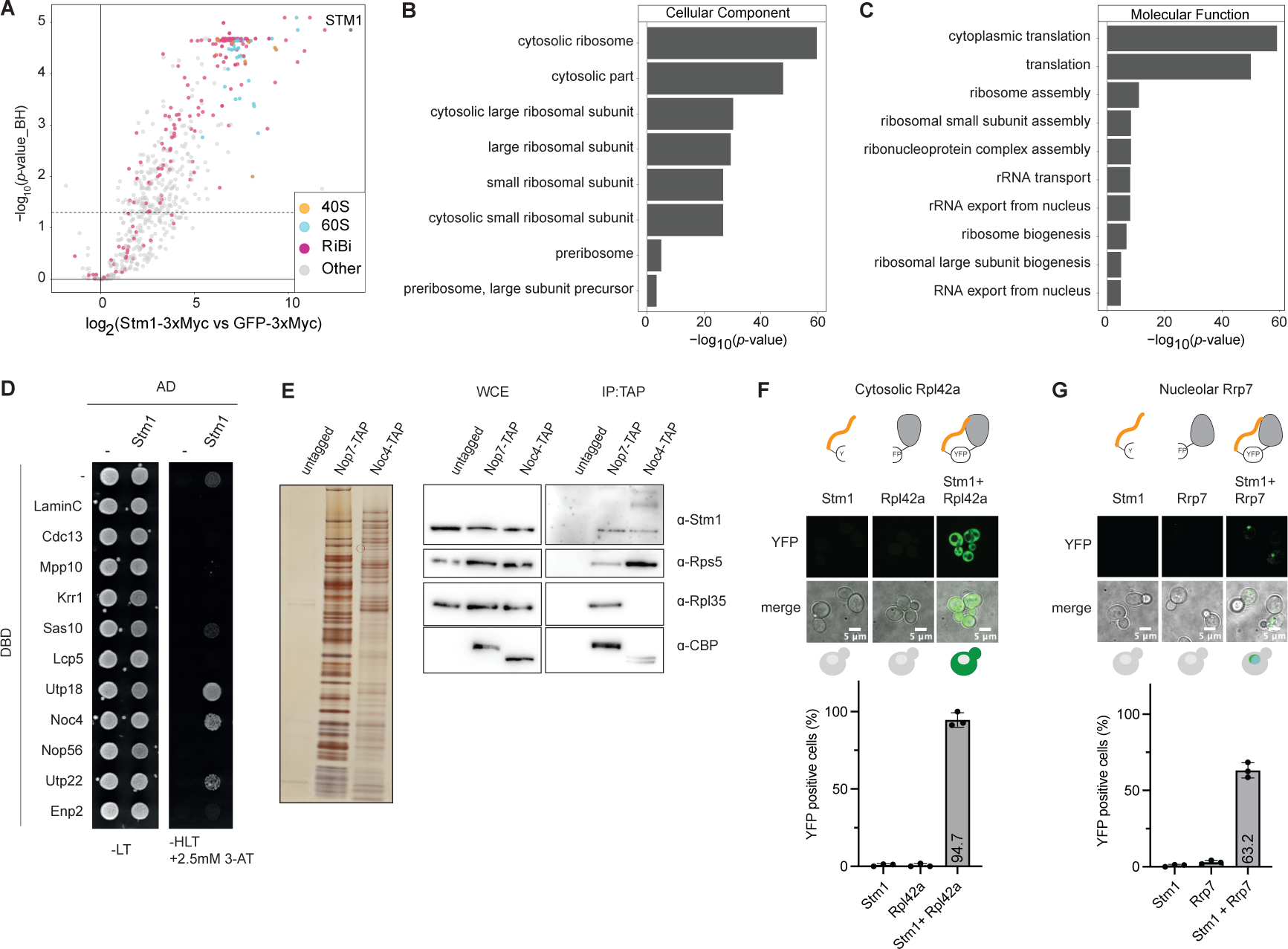
Stm1 interacts with pre-ribosomal particles. **A**) Volcano plot showing the Stm1 interactome as log_2_ fold enrichment of proteins co-purified with Stm1-3xMyc versus GFP-3xMyc controls. The significance was calculated by two-sided unpaired Student’s t-tests, *p*-value was corrected for multiple hypothesis using the Benjamin-Hochberg method. N=3 independent replicates. 40S (orange), 60S (blue) ribosomal proteins and RiBi factors (purple) are depicted. **B)** and **C**) Gene ontology enrichment for Cellular Components (**B**) and Molecular Function **(C)** was calculated for Stm1 interactors (defined as log_2_FC> 2 and *p-*value< 0.05). The significant of the enrichment is reported as the -log_10_ of *p*-value. **D)** Two-hybrid analysis of Stm1 fused to the GAL4 activation domain (AD) or AD control (-) with SSU components or various controls (-, Lamin C, Cdc13) fused to GAL4 DNA binding domain (DBD). Colonies were grown for 4 days at 30^°^C on permissive auxotrophic SD -Leu, -Trp (-LT), or selective -His, -Leu, -Trp agar media, supplemented with 2.5mM 3-amino-1,2,4-triazole (-HLT +2.5mM 3-AT). Representative image from N=3 independent replicates. Linked to **Sup 4A.** **E)** Tandem affinity purification (TAP) of untagged, Nop7-TAP or Noc4-TAP pre-ribosomal particles. Silver stained SDS-PAGE (left panel) and western blot (right panel) are shown. Blots were probed as indicated with antibodies to CBP (Nop7-TAP and Noc4-TAP), Stm1, Rps5, Rpl35. Representative gel from N=3 independent replicates. **F)** and **G**) Schematic representation of BiFC assay. Estradiol inducible C-terminal YFP fragment Vc is fused to Stm1 (orange), and constitutive N-terminal YFP fragment Vn fragment is cloned in frame with Rpl42a (**F**) or Rrp7 (**G**). If Stm1 interacts with Rpl42a or Rrp7, the Vc and Vn fragments reconstitute YFP and thus fluorescence (top panel). Representative images of estradiol induced (10nM for 2h) cells expressing Vc-Stm1 (Stm1) or Rpl42a-Vn (Rpl42a) or Vc-Stm1 and Rpl24a-Vn (Stm1 + Rpl42a) are shown in the middle panel. The BiFC assays were quantified and shown as percentage of YFP positive cells (bottom bar graph). At least 50 cells per sample and independent replicate was analyzed (N=3). Data values are shown as dots as well as mean and standard deviation.

To dissect the subcellular localization of Stm1, we imaged N- and C-terminally GFP-tagged Stm1 and, as expected, found that GFP-Stm1 was predominantly cytosolic (**Sup 2E**). To visualize its potential nucleolar localization, we implemented bimolecular fluorescence complementation (BiFC) assays (41). Briefly, Stm1 was fused to the C-terminal half of YFP (Vc), and the N-terminal YFP half (Vn) to the nucleolar protein Rrp7 (Rrp7-Vn), a component of the SSU sub-complex UtpC (42). For control, the Vn-fragment was fused to the cytosolic ribosome subunit Rpl42a (Rpl42a-Vn), which in the ribosome crystal structure (30) binds adjacent to Stm1 (**Fig 2F and G**). As expected, Stm1-Vc binds Rpl42a-Vn in the cytoplasm, while no YFP signal was observed in cells expressing either fusion protein alone, demonstrating specificity of the assay. Importantly, Stm1-Vc readily interacted with Rrp7-Vn, as visualized by the nucleolar YFP fluorescence. Quantification of these BiFC assays revealed that over 60% of cells expressing both Stm1 and Rrp7 fusions gained a nucleolar YFP signal, while over 90% stained positive for the Rpl42a cytoplasmic control (**Fig 2F and G**). Taken together, these results demonstrate that Stm1 exhibits a versatile behavior in the cell, partitioning between the mature ribosomes in the cytoplasm and RiBi and pre-RPs in the nucleolus.

### Stm1 controls the transcription of RPG targets of Ifh1

To stage Stm1 function during early steps of ribosome biogenesis, we analyzed the synthetic genetic analysis database for negative genetic interactions (43). Interestingly, several components regulating the RNA Pol II machinery such as Kin28 or subunits of the mediator complex (Med) were synthetically sick with *stm1Δ* (**Sup 3A**). Furthermore, the slow RNA Pol II mutants *rpo21/rpb1-N479S* and *rpo21/rpb1-N1082S* (44) exhibit negative genetic interaction with *stm1Δ* (**Sup 3B**), suggesting that Stm1 may be essential in cells with reduced RNA Pol II activity. To confirm this hypothesis, we engineered strains with the anchor-away system (45) carrying two copies of the *RPO21* gene: an untagged copy of wild-type *RPO21* or the slow mutants *rpo21-N479S*, *rpo21-N1082S and rpo21-H1085Q* (44), and FRB-tagged *RPO21*, which after expression is sequestered in the cytoplasm upon addition of the dimerization compound. This genetic assay confirmed that the slow RNA Pol II mutants *RPO21-N479S*, *N1082S* and *H1085Q* exhibit a synthetically lethal interaction when combined with *stm1Δ* (**Fig 3A**), demonstrating that Stm1 is needed when RNA Pol II activity is perturbed.

**Figure 3:**
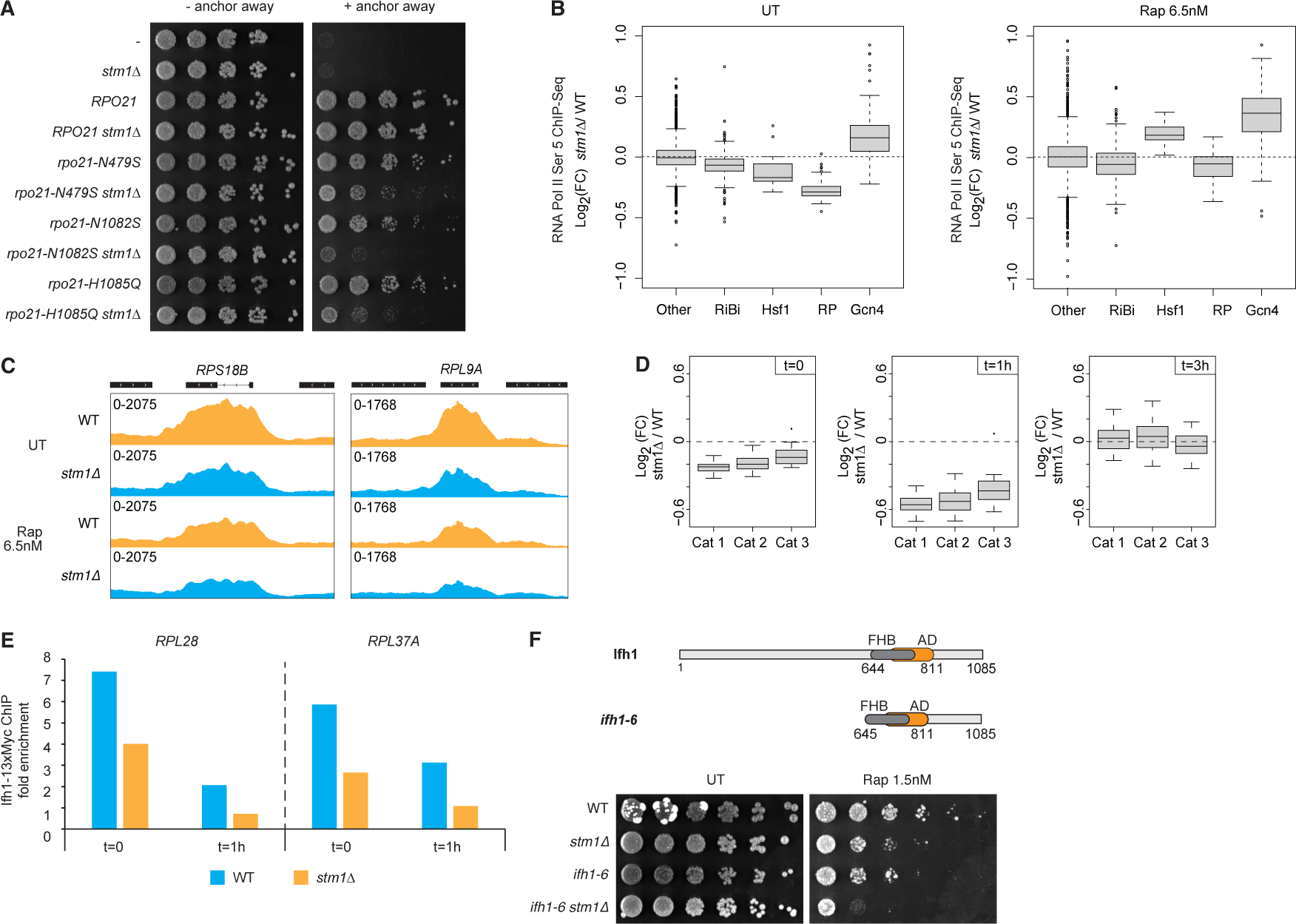
Stm1 promotes Ifh1 activity at RPGs. **A)** Serial dilution spotting assay to monitor growth on YPD media of WT (-) or anchor away strains (45). Anchor away strains were either WT or deleted for *STM1* (*stm1Δ*) and carry an extra copy of genomically-integrated RPO21 WT (*RPO21*) or different point mutations (*rpo21-N479S, rpo21-N1082S, rpo21-N1085Q).* Nuclear depletion of endogenous *RPO21* (anchor-away) was initiated by the addition solvent control (-anchor away) or 1µM rapamycin (+anchor away). Cells were incubated for 3 days. Representative image of N=3 independent replicates. **B)** Box plots showing RNA Pol II Ser 5 occupancy. The differential analysis of *stm1Δ* versus WT cells is shown for ribosomal protein genes (RP (n=138)), ribosome biogenesis genes (RiBi (n=566)), Gcn4 targets genes (Gcn4 (n=108)), heat shock genes (Hsf1 (n=18)) or others (n=4311) genes. Data are displayed as the log_2_ normalized fold change of RNA Pol II S5 occupancy in *stm1Δ* versus WT cells, on the left for untreated conditions (UT) and on the right 3 hours after low dose rapamycin treatment (Rap 6.5nM). The independent replicate is shown in **Sup 3D**. The signal was quantified for each RNA Pol II transcribed gene between the transcription start site (TSS) and transcription termination site (TTS). The category of genes was defined according to scientific literature: (56) for RPG, RiBi, (86) for Hsf1, (47) for Gcn4 targets. **C)** Genome browser tracks showing RNA Pol II S5 ChIP-Seq read counts at open reading frames of the *RPS18B* (left panel) and *RPL9A* (right panel) genes in WT (orange) and *stm1Δ* (blue) cells, treated (Rap 6.5nM) or not (UT) with rapamycin for 1 hour. **D)** Box plots comparing differential RNA Pol II Ser 5 occupancy of ribosomal protein genes (RP (n=138)) categories number 1 (Cat 1), number 2 (Cat 2) and number 3 (Cat 3) (56) in WT and *stm1D* strains. Data are displayed as the log_2_ normalized fold change of RNA Pol II Ser 5 occupancy in *stm1Δ* versus WT cells, treated or not (t=0, left panel) with 6.5nM rapamycin for 1h (t=1, central panel) or 3h (t=3, right panel). An independent replicate is shown in **Sup 3I.** **E)** Bar plot showing Ifh1-13xMyc occupancy fold enrichment at the RPGs *RPL28* (left panel) and *RPL37A* (right panel) analyzed by ChIP-qPCR and normalized on *ACT1.* WT and *stm1Δ* cells were treated or not (t=0) for 1h with 6.5nM rapamycin (t=1h). An independent replicate is shown in **Sup 3J.** **F)** Schematic of ifh1-6 and ifh1 (top) used in growth assays (bottom). Serial dilution spotting assay to compare growth of WT, *stm1Δ*, the Ifh1 truncation mutant *ifh1-6* (AA 645-1085), and the *ifh1-6 stm1Δ* double mutant on YPD media. Cells were either treated or not (UT) with 1.5nM rapamycin (Rap 1.5nM) and incubated for 3-4 days. A representative image from N=3 independent experiments is shown.

While Stm1 associates with pre-ribosomal complexes, Stm1 was not present in pull-downs of Rpb8-TAP, a common component of RNA Pol I, Pol II and Pol III complexes (36) (**Sup 3C**), excluding the possibility that Stm1 stably interacts with the transcription machinery. To probe whether Stm1 regulates expression of a specific gene cluster, we next measured the occupancy of RNA Pol II Ser 5 by chromatin immunoprecipitation sequencing analysis (ChIP-Seq) in wild-type and *stm1Δ* cells before and after mild rapamycin treatment (**Fig 3B and C, Sup 3D-H, Supplementary table 3**). Interestingly, we found a decrease in occupancy of RNA Pol II at RPGs in *stm1Δ* cells, whereas the reduction was notably less pronounced at RiBi genes. Moreover, *stm1Δ* upregulates Gcn4 targets in absence and presence of low dose rapamycin, and Hsf1 targets only upon TORC1 inhibition, indicating a general level of cellular stress (46, 47) (**Fig 3B, Sup 3D-H**). Gcn4 upregulation did not depend on eIF2alpha/Sui2 phosphorylation (48) (**Sup 1J**) and might thus originate from alternative pathways that impact ribosome biogenesis (49–52). While transcription of RiBi genes is specifically regulated by Sfp1 (53–55), RPGs are mainly targeted by Ifh1 (56). RPGs are subclassified in three categories, two of them mainly activated by Ifh1 and the third one mostly regulated by Sfp1 (56). Interestingly, detailed analysis revealed that in *stm1Δ* cells, the RNA Pol II occupancy at the three RPGs categories is downregulated in the absence and presence of low dose of rapamycin (**Fig 3D, Sup 3I, Supplementary table 3**). Since there is no prominent effect of *stm1Δ* on RNA Pol II occupancy at Sfp1 targets of the RiBi genes, we hypothesized Stm1 predominantly promotes Ifh1 activity.

To test this hypothesis, we measured the occupancy of Ifh1-13xMyc by ChIP-qPCR in wild-type and *stm1Δ* cells. We observed that Stm1 promotes Ifh1 binding at RPG promoters of *RPL28* and *RPL37A* relative to *ACT1* in both untreated and rapamycin treated conditions (**Fig 3E, Sup 3J**). To ensure Stm1 effect is specific for Ifh1 targets, the *RPL18A* endogenous promoter was replaced by the doxycycline repressible TetO7 promoter. Stm1 knock out did not affect the growth defect of repressible *RPL18A* upon addition of doxycycline suggesting that Stm1 absence does not further perturb the transcription of RPGs when not activated by Ifh1 (**Sup 3K**). Taken together, these data suggest that Stm1 enhances transcriptional regulation of RPGs, by directly or indirectly promoting Ifh1 activity.

To genetically validate that Stm1 functions as an Ifh1 activator *in vivo*, we deleted *STM1* in strains with altered Ifh1 activity. The truncation mutant *ifh1-6,* encompassing the Forkhead binding domain (FHB), activation domain (AD) and C-terminus of Ifh1, stably associates with RPG promoters (12). Interestingly, deletion of *STM1* and exposure to low dose rapamycin compromised viability of *ifh1-6* cells, indicating that Stm1 interferes with altered Ifh1 dynamics (**Fig 3F**).

### The Stm1 C-terminal IDR confers rapamycin sensitivity independent of its hibernation function

The crystal structure of Stm1 bound to ribosomes indicates the N-terminus binds the 60S particle and the linker region bridges the 40S and 60S ribosomal subunits. However, the Stm1 structure has only been resolved with its central domain, while the C-terminus is predicted to be unstructured and has not been characterized (30, 57).

To test the functional relevance of the N- and C-terminal Stm1 domains (**Fig 4A**), we expressed 3xMyc-tagged wild-type or truncation mutants, stably integrated as single copy in *stm1Δ* strains. Western blot analysis confirmed that Stm1 variants are expressed at the expected size with comparable steady-state levels (**Fig 4B, lanes 1-6**). Growth assays revealed that rapamycin-treated cells tolerate short truncation of the Stm1 N-terminus (*stm1ΔN)*, although this domain is thought to mediate the Stm1 hibernation function. In contrast, cells expressing Stm1 lacking the C-terminus (*stm1ΔC)* displayed inhibited growth on rapamycin-containing media, like *stm1Δ* controls (**Fig 4C**). Strikingly, overexpression of a GFP-tagged C-terminal Stm1 domain (GFP-C) was able to rescue the rapamycin-induced growth defect (**Fig 4D, E**) suggesting that Stm1’s C-terminal domain is necessary and sufficient for its starvation function *in vivo*.

**Figure 4:**
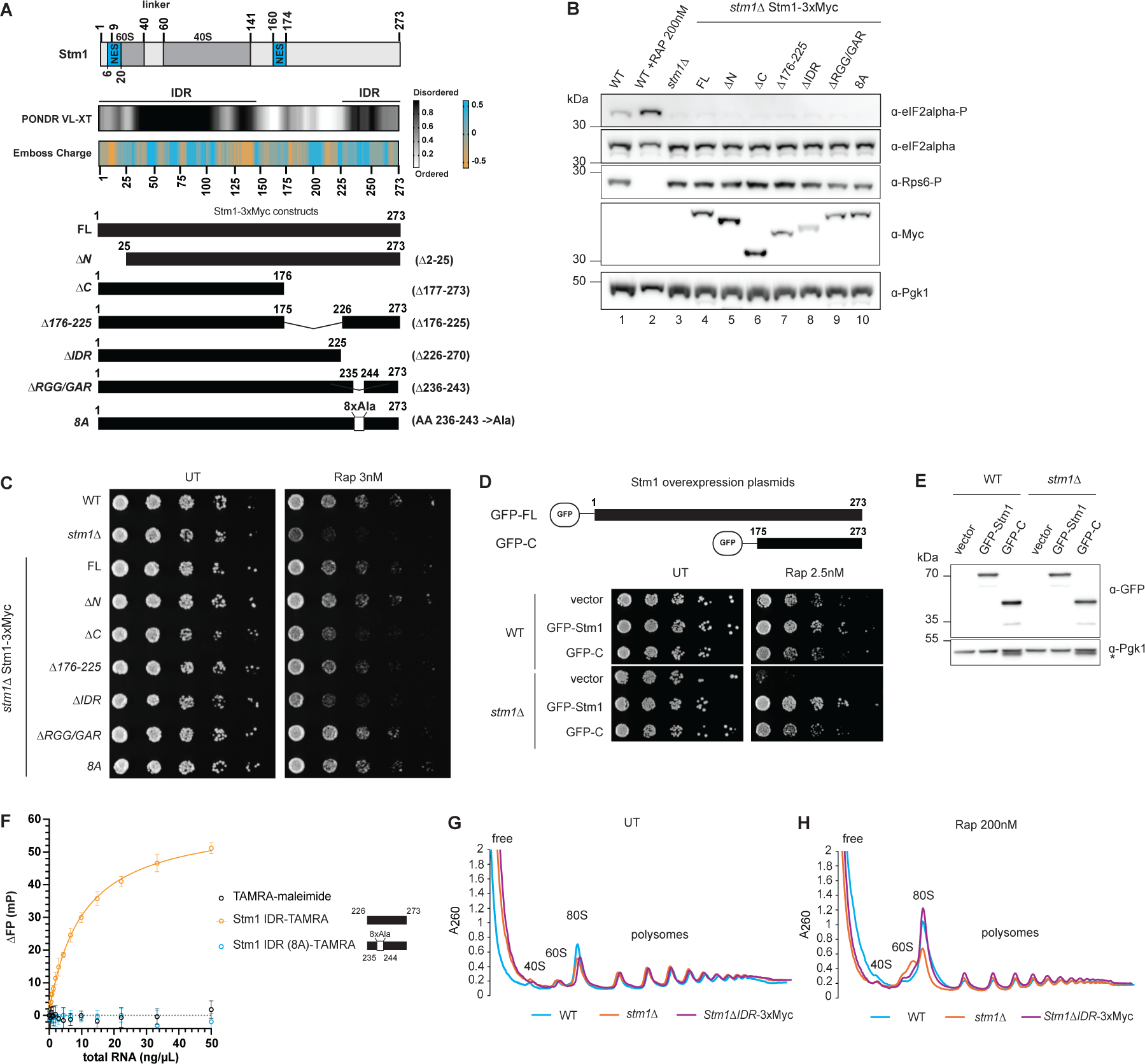
Stm1 C-terminal IDR drives rapamycin sensitivity. **A)** Schematic drawing with amino acid numbers of Stm1 depicting the 60S and 40S binding domains separated by a linker region (30), and two predicted nuclear export signals (NES) (64) (top). The disorder score (0–1 PONDR VL-XT and amino acid charge (-0.5 to +0.5) using Emboss Charge (middle) is shown below. The C-terminally tagged full length Stm1-3xMyc (FL) and different mutants (*ΔN*, *ΔC*, *Δ176-225*, *ΔIDR*, *ΔRGG/GAR*, *8A*) used in this study are illustrated, with the relevant amino-acid numbers indicated. They were cloned into single integration pSIV vectors (40) and integrated at the *URA3* locus. **B)** Western blot of total lysates prepared from WT or *stm1Δ* cells treated or not with 200nM rapamycin for 2 hours (Rap 200nM). Stm1-3xMyc and the indicated Stm1 mutants were integrated in *stm1D* cells and included in the analysis. Blots were probed with antibodies against phospho-eIF2alpha (Sui2 Ser51), eIF2alpha, phospho-Rps6 (Ser232/233), Myc, and Pgk1 to control equal loading. A representative image of N=3 independent replicates is shown. **C)** Serial dilution spotting assays to monitor growth on YPD media of WT, *stm1Δ*, and *stm1Δ* strains stably expressing 3xMyc-tagged full length (FL) or the indicated mutants (*ΔN, ΔC, Δ176-225, ΔIDR, ΔRGG/GAR, 8A*). Cells were either treated or not (UT) with 3nM rapamycin (Rap 3nM) and incubated for 2 days. A representative image of at least 3 independent experiments is shown. **D)** Serial dilution spotting assay to monitor growth of WT and *stm1Δ* cells harboring an empty control plasmid (vector), or plasmids overexpressing N-terminal GFP-tagged full length Stm1 (GFP-Stm1) or its GFP-tagged C-terminal domain (GFP-C, AA 175 -273). The constructs are schematically indicated on top. Cells were either treated or not (UT) with 2.5 rapamycin (Rap 2.5nM) and incubated for 3 days. A representative image from N=2 independent experiments is shown. **E)** Western blot confirming expression of GFP-Stm1 and GFP-C in WT or *stm1Δ* cells. Cells harboring an empty plasmid (vector) are used for control. Blots were probed with antibodies against GFP and endogenous Pgk1 to control equal loading. The asterisk (*) marks the signal from the GFP antibody migrating close to Pgk1. A representative image of N=2 independent replicates is shown. **F)** Fluorescent polarization assay to test binding of yeast total RNA (total RNA (ng/µL)) to TAMRA-labelled Stm1-IDR wild type peptide (AA 226-270) (Stm1 IDR-TAMRA) or a corresponding mutant peptide (Stm1 IDR (8A)-TAMRA) where the arginine and lysine residues in the RGG/GAR motifs were changed to alanine (AA 236-243 ->8A). TAMRA-maleimide controls non-specific binding. N=3 independent replicates. Note that Stm1 binds RNA through its RGG/GAR motifs. **G)** and **H)** Ribosome and polysome profiling of WT (blue), *stm1Δ* (orange) and *stm1ΔIDR*-3xMyc (purple) cells recorded at A_260_ absorbance in treated (panel **H**) or not (UT, panel **G**) with 200nM rapamycin for 1 hour (Rap 200nM). The position of free, 40S, 60S, 80S and polysomes is indicted on top.

To gain mechanistic insights how Stm1 may promote Ifh1 activity, we predicted functionally relevant domains (58, 59) (**Fig 4A**). We identified putative nuclear export signals (NES) and IDR domains at the N- and C-termini, and the C-terminal IDR encompasses RGG and GAR RNA binding-motifs (**Fig 4A**). Consistent with previous reports about the ability of Stm1 to bind nucleic acids (60–62), fluorescence polarization (FP) assays revealed that TAMRA-labeled Stm1-IDR readily bound total RNA, only in the presence of the RGG/GAR motif (236-243 region of Stm1) (**Fig 4F**).

To test the functional relevance of these Stm1 domains, we constructed mutants of the C terminal region of Stm1: *Δ176-225*, *ΔIDR (Δ226-270)*, *ΔRGG/GAR (Δ236-243)*, *8A (AA236-243 ->Ala)* (**Fig 4A**). All mutants were stably integrated as single copy, tagged with 3xMyc in *stm1Δ* cells and were expressed at the expected size. We excluded global impairment of protein translation or TORC1 activity based on Sui2 and Rps6 phosphorylation levels. (18, 63) (**lanes 7 to 10 in Fig 4B**). Importantly, cells expressing *stm1ΔIDR* are sensitive to rapamycin comparable to *stm1Δ* control cells, while *stm1Δ176-225* cells did not display inhibited growth. In contrast, rapamycin-sensitivity was not dependent on a functional RGG/GAR motif (*stm1ΔRGG/GAR*), suggesting that RNA-binding by these motifs is not critical for Stm1 function (**Fig 4C**). We then focused on Stm1ΔIDR-3xMyc and analyzed ribosome profiles by sucrose gradients separation in untreated and rapamycin-treated conditions. As expected, truncation of the C-terminal IDR did not impact 80S levels (**Fig 4G and H**). Taken together, we conclude that the C-terminal IDR domain of Stm1 is necessary and sufficient for its growth-promoting function, which is independent from ribosome protection.

### Stm1 IDR drives the interaction with RiBi and CURI complex

To test if previously identified Stm1 interactors bind through its C-terminus, we first used two-hybrid analysis with *stm1ΔN* and *stm1ΔC* mutants. While some components such as Krr1 or Enp2 preferentially bind to *stm1ΔC*, most nucleolar interactors require the presence of the unstructured C-terminus, and their interaction seems greatly increased in *stm1ΔN* mutants (**Sup 4A**). Furthermore, we conducted a comparison between the interactomes derived from the AP-MS of Stm1-3xMyc and Stm1ΔIDR-3xMyc. Following normalization for the abundance of purified Stm1, we observed a remarkable reduction in interactions with ribosomal proteins, translation regulators and RiBi in the Stm1ΔIDR-3xMyc pulldown, underscoring the critical relevance of the C-terminal IDR region (**Fig 5A, 5B, Sup 2A, B, C, Supplementary table 2**). In stark contrast, binding of RiBi to Stm1ΔN-3xMyc mutant exhibited an increase compared to full length Stm1-3xMyc and Stm1ΔIDR-3xMyc (**Sup 4B-G, Supplementary table 4**), possibly due to nuclear retention as the protein lacks a predicted nuclear export signal (NES) at amino acids 6-20 (**Fig 4A**) (64). We conclude that Stm1ΔN and Stm1ΔIDR have distinct protein networks, highlighting separable functions of Stm1.

**Figure 5:**
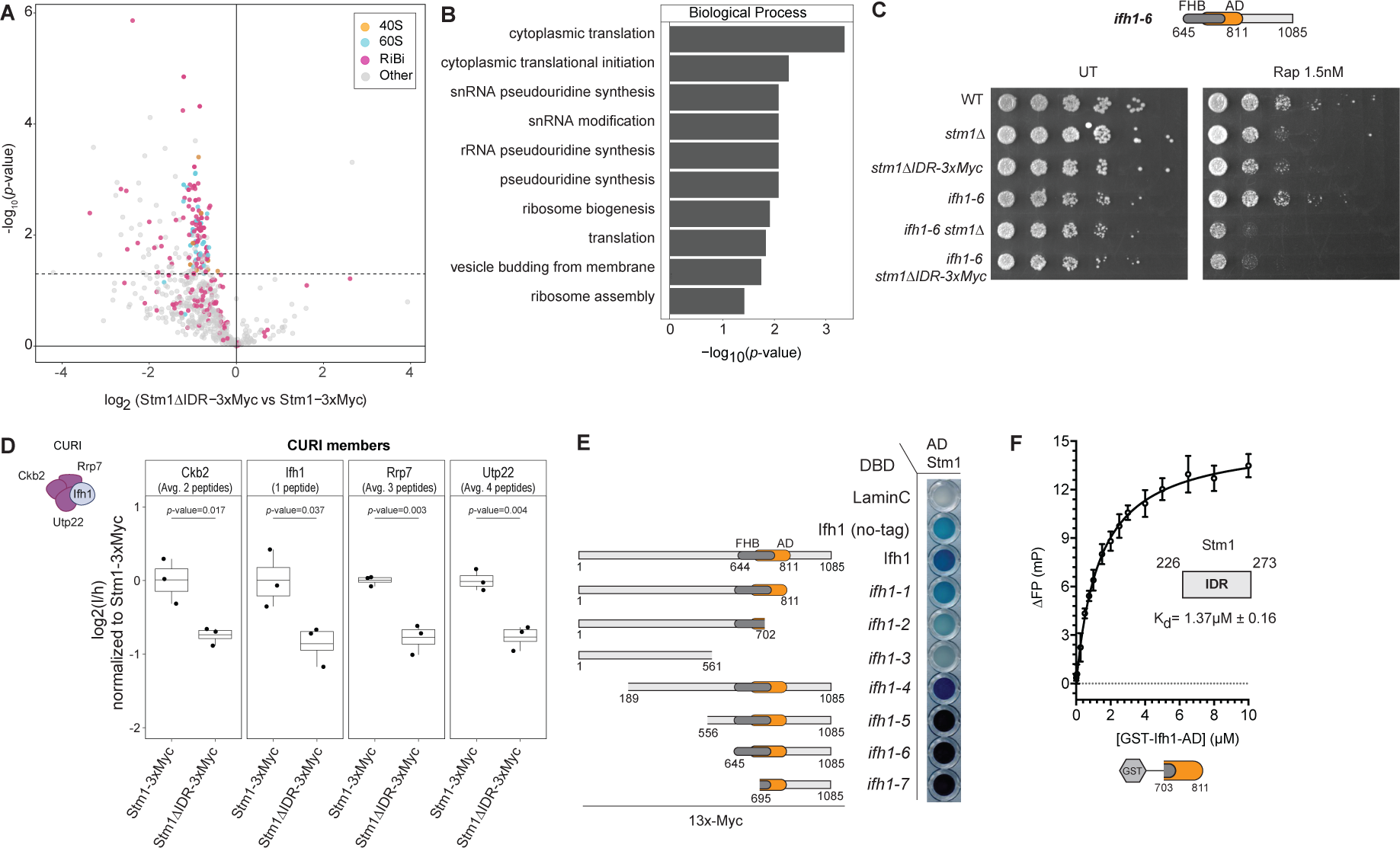
The Stm1-IDR drives interaction with CURI subunits and promotes Ifh1 activity. **A**) Volcano plot showing the log_2_ fold change enrichment for proteins co-purified with 3xMyc-tagged Stm1ΔIDR in comparison to Stm1. The protein abundance was normalized to Stm1 levels after AP-MS. The significance is calculated from two-sided unpaired Student’s t-test. N=3 independent replicates. Proteins of the 40S (orange) and 60S (blue) ribosomal subunits and RiBi factors (purple) are highlighted. Linked to **Sup 2A, B, C.** **B**) The gene ontology for Biological Process was calculated for the Stm1 interactors identified in **Fig 5A** (defined as log_2_FC< -1 and *p-*value< 0.05). The significance of the enrichment is reported as the -log_10_ of *p*-value. **C**) Schematic representation of the *ifh1-6* truncation mutant encompassing amino acids 645 to 1085 including its forkhead binding domain (FHB) and the activation domain (AD) (top). Serial dilution spotting assay (bottom) comparing growth of WT, *stm1Δ*, *stm1ΔIDR-3xMyc, ifh1-6* and *ifh1-6 stm1Δ* and *ifh1-6 stm1ΔIDR-3xMyc* double mutants on YPD media. Cells were treated or not (UT) with 1.5nM rapamycin (Rap 1.5nM) and incubated for 3-4 days N=3 independent replicates. **D**) Targeted PRM measurement of Ifh1 and CURI subunits co-purified with Stm1. Box plots showing the log_2_ fold change enrichment of Ckb2, Ifh1, Rrp7 and Utp22 in Stm1ΔIDR -3xMyc compared to Stm1 WT (Stm1-3xMyc). Data values (normalized for Stm1 abundance) are shown as log_2_ ratio of light (endogenous peptide) to heavy (stable heavy isotope-labeled peptide) (l/h). Protein abundance is calculated from the average intensity of the measured peptides (all measured peptides are reported in **Sup 5A**). The number of peptides used for each protein is indicated below the protein name. Independent replicates are represented by a dot (N=3), with the box plot boundaries indicating the quantiles Q1 (25%) and Q3 (75%). The average value is indicated by a line across the box. Statistical analysis was performed with Student t-test’s, two sided unpaired. **E**) β-galactosidase assay of two-hybrid pairs to test interactions of Stm1 fused to the GAL4 activation domain (AD Stm1) and previously described Ifh1 truncation mutants (12) fused to the GAL4 DNA binding domain (DBD). Lamin C and untagged Ifh1 were used as negative and positive controls, respectively. The plate was imaged after 19 hours. A representative image from N=3 independent replicates is shown. **F**) Fluorescent polarization assay quantifying binding of the GST-tagged Ifh1 activation domain (AA 703-811) (GST-Ifh1-AD) and TAMRA-labelled Stm1-IDR peptide (AA 226-273) (Stm1 IDR-TAMRA), normalized to GST. The dissociation constant (K_d_) is indicated. N=3 independent replicates.

The RiBi factors Utp22 and Rrp7, and the CK2 subunit Ckb2 bind Ifh1 (CURI complex) in the nucleolus, thereby sequestering Ifh1 and consequently downregulating the expression level of RPGs (11, 12). Since Stm1 controls RPG transcription (**Fig 3B, Sup 3D**) and interacts with nucleolar and nuclear RiBi factors (**Fig 2A**), we hypothesized that Stm1 may bind Ifh1 through its IDR. Indeed, the genetic combination of *stm1ΔIDR* with the *ifh1-6* truncation mutant is detrimental for cell growth (**Fig 5C**). Two-hybrid results (**Sup 4A, H**) and quantitative targeted parallel reaction monitoring (PRM)-MS analysis (65) showed that Stm1 binds Ifh1 and all subunits of the CURI complex (Ckb2, Utp22, and Rrp7) with at least an enrichment of 5-fold compared to GFP-3xMyc control (**Sup 4I-K, Sup 5A, Supplementary table 5**). Importantly the Stm1 IDR domain is necessary to efficiently interact with Ifh1 and the CURI subunits (**Fig 5D, Sup 4I-K, Supplementary table 5**), in contrast to AA176-225 (**Sup 4H**). We conclude that the growth promoting function of Stm1 requires its C-terminal IDR, which is necessary for binding to Ifh1 and members of the CURI complex.

To disentangle the mechanism of interaction between Stm1 and Ifh1, we combined different Ifh1 truncation mutants (12) with Stm1 in two-hybrid assays. While the amino-terminal domain (AA 1 – 702, *ifh1-2*) was unable to bind, a fragment encompassing the AD and parts of the C-terminal FHB domain (*ifh-*7) was sufficient to interact with full-length Stm1 (**Fig 5E**). To validate whether these two domains can directly interact *in vitro*, we expressed and purified the GST-tagged AD-domain of Ifh1 (GST-Ifh1-AD, amino acids 703-811) and used FP to measure its binding to a peptide encompassing the IDR domain of Stm1. Interestingly, Ifh1-AD directly bound Stm1-IDR, with a K_d_ of approximately 1.4 µM (**Fig 5F**). Taken together, we conclude that Stm1 interacts directly through its C-terminal IDR with the activation domain of Ifh1, and that this interaction is required for Ifh1-mediated cell growth and RPG activity.

## Discussion

Here we report that ribosome dormancy factor Stm1 promotes ribosome biogenesis (**Fig 6A, B**). In unchallenged conditions, Stm1 localizes to the nucleolus where it engages with pre-ribosomal particles and members of the CURI complex, including the RPG transcription factor Ifh1. Stm1 has a C-terminal IDR domain that confers resistance to TORC1 inhibition and is required for binding to RiBi and CURI subunits. Moreover, the IDR of Stm1 directly binds to the activation domain of Ifh1, thereby regulating cell growth by enhancing RPG transcription.

**Figure 6:**
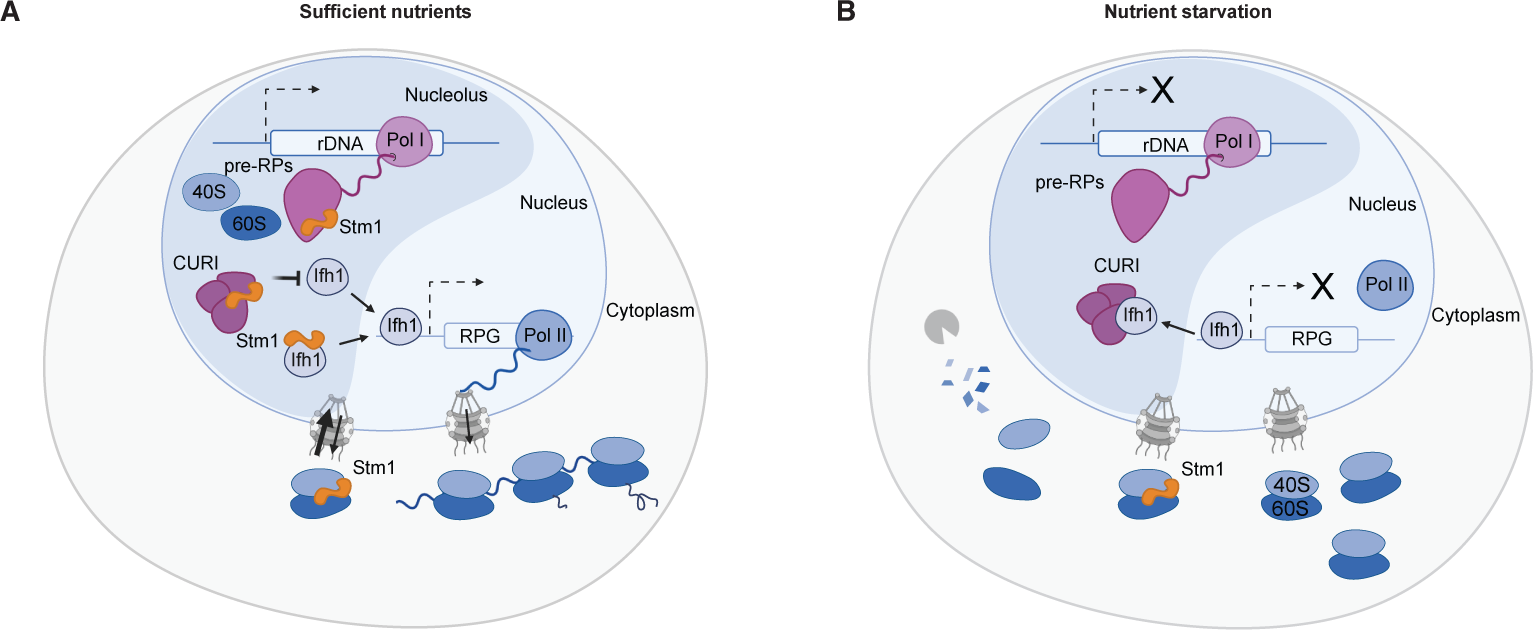
The hibernation factor Stm1 protects 80S ribosomes upon starvation but promotes RPG transcription by activating Ifh1 during exponential growth. **A)** and **B**): Model depicting Stm1 functions during exponential growth (**A**) and nutrient starvation (**B**). During starvation conditions, ribosome biogenesis is downregulated and cytoplasmic Stm1 protects a subset of ribosomes from degradation by preventing dissociation of the 40 and 60S ribosomal subunits (right panel). During nutrient rich conditions, Stm1 localizes to the nucleolus and interacts with CURI components and the transcription factor Ifh1. Stm1 directly binds Ifh1 and may help releasing Ifh1 from the CURI, thereby increasing RPG transcription. Stm1 also interacts with pre-ribosomes (pre-RPs), and my thus promote different steps of ribosome maturations (left panel). Together, these results suggest that Stm1 regulates ribosome homeostasis, and links ribosome production and degradation.

### Stm1 promotes RiBi transcription by regulating Ifh1 activity

Previous studies described Stm1 as a ribosome dormancy factor and TORC1-regulated translation inhibitor. These functions involve the Stm1 N-terminal domain that intercalates in the ribosome mRNA tunnel and the linker region (Stm1 AA40-60) that bridges the 40S and 60S Stm1 binding domains (**Fig 4A**). Indeed, *stm1Δ* cells or cells expressing an N-terminal truncation (Δ2-40 or Δ41-80) show reduced 80S ribosomes after starvation (24). In this study, we used a shorter N-terminal truncation of Stm1 (*stm1ΔN*: Δ2-25) that does not affect the linker region. Surprisingly, *stm1ΔN* mutant co-purifies predominantly with RiBi factors and cells are not sensitive to rapamycin, indicating that the ribosome protection function of Stm1 does not explain its TORC1 inhibition phenotype. Instead, our results demonstrate that Stm1 plays a second, rapamycin-sensitive function by promoting ribosome biogenesis in nutrient-rich conditions. Moreover, the *stm1ΔN* mutant may delay an intermediate and transient state of the Stm1 cycle across cellular compartments. Surprisingly, while the concentration of cytoplasmic ribosomes is decreased in *stm1Δ* cells, ribosome degradation was comparable to wild-type controls (**Fig 1C-I, Sup 1D-H, K, L**). Several lines of evidence suggest that Stm1 promotes ribosome biogenesis by increasing nuclear Ifh1 activity. First, *stm1Δ* cells display a drop in 35S rRNA levels as a possible consequence of lower expression of RNA Pol I, II and III subunit Rpb5 (**Fig 1E-G, Sup 1F**). Second, Stm1 is essential for cell growth when RNA Pol II (**Fig 3A**) or Ifh1 activity are reduced or altered (**Fig 3F, G**). Third, binding of RNA Pol II and Ifh1 to RPG promoters is reduced in the absence of Stm1 (**Fig 3E, Sup 3J**). Significantly, resistance to rapamycin requires Stm1’s C-terminal IDR, which directly binds Ifh1 and members of the CURI complex. The exact mechanism by which Stm1 modulates Ifh1 activity remains elusive. Stm1 binding to the activation domain of Ifh1 may increase its ability to recruit Pol II, concomitant with rRNA synthesis and processing (11, 12). This interaction may be transient, and structural work will be required to unravel the function of Stm1 when bound to Ifh1. Alternatively, but not mutually exclusive, Stm1 and Ifh1 may compete for binding to the nucleolar CURI complex comprised of Utp22 and Rrp7, and the CK2 kinase subunit Ckb2, and thus Stm1 may help to release Ifh1 to allow its binding to RPG promoters. Finally, Stm1 was identified as a suppressor of Tom1 (66), which is required to prevent aggregation of ribosomal proteins (9, 67, 68). Stm1 may thus impact Ifh1 solubility in the crowded nuclear/nucleolar compartments. Irrespective of the molecular mechanism, the Stm1 IDR region contributes to the nuclear/nucleolar interaction network and integrate environmental cues towards Ifh1 and other interactors (69, 70). Further studies will be required to uncover how Stm1 modulates Ifh1 activity.

### Stm1 may promote ribosome biogenesis independently of rRNA transcription and RiBi transcription

While Stm1 promotes Ifh1 activity, it also promotes ribosome biogenesis in the nucleolus downstream of rRNA transcription. Indeed, Stm1 co-purifies with pre-ribosomal particles (**Fig 2A-E)**, and its association is greatly increased in Stm1 mutants lacking an intact N-terminus (*stm1ΔN*), most likely because this domain contains a nuclear export signal (**Sup 4A-G**). Interestingly, Stm1 interacts with several components of the SSU processome including Noc4, Mpp10, Krr1 and Utp18 (**Fig 2D and E, Sup 4A**). The SSU processome associates with the nascent pre-rRNA and works in concert to orchestrate RNA folding and cleavage, and targets pre-ribosomal rRNA for degradation by the RNA exosome. Utp18 is crucial for interaction with exosome and TRAMP complexes and the maturation of the SSU processome (71). Interestingly, Stm1 directly binds RNA, in part through a conserved RGG-motif in its C-terminus (**Fig 4F**). While the RNA-binding motif is not essential to confer rapamycin-sensitivity *in vivo* (**Fig 4C**), Stm1 binds G4 quadruplex and purine triplex motifs, RNA folds which are present in rRNA and telomeric structures (62). Finally, Stm1 was also identified as RNA binding protein in response to stress (72) and as a component of the cytoplasmic mRNA degradation machinery, connecting mRNA decapping to ribosome function. Indeed, Stm1 promotes mRNA de-adenylation and decay of a subclass of mRNAs, including the Dhh1 mRNA targets EDC1 and COX17 (73). Thus, Stm1 plays multiple roles in the nucleus and cytoplasm relating to the regulation of RNA-mediated processes. Although the function of Stm1 in pre-90S pre-ribosomal particles remains to be explored, we speculate that it may alter rRNA processing by regulating SSU activity.

### Stm1 links protein translation and ribosome biogenesis

In exponentially growing cells, Stm1 specifically binds 80S ribosomes and polysomes in a salt-dependent manner (24, 74) and inhibits translation (34, 75). A fraction of nuclear/nucleolar Stm1 enhances Ifh1-dependent RPG transcription and promotes ribosome biogenesis. Nutrient starvation inactivates TORC1 and inhibits direct phosphorylation of Stm1. Dephosphorylated Stm1 clamps the two ribosomal subunits, preventing their dissociation (76, 77), which inhibits ribosome degradation by autophagy and UPS (24). Thus, Stm1 not only controls ribosome production, but also ribosome hibernation in response to nutrient starvation. Interestingly, these two activities are mutationally separable, as they are mediated through distinct Stm1 domains. The C-terminus of Stm1 contains an unstructured IDR, which mediates binding to Ifh1 and the nuclear and nucleolar RiBi. In contrast, Stm1 lacking this IDR domain maintains its ribosome hibernation activity (**Fig 4H**), and surprisingly, confers rapamycin sensitivity (**Fig 4C**). Indeed, the reduced number of ribosomes in *stm1Δ* cells is not caused by increased degradation, but rather decreased ribosome production. It will be interesting to examine whether other ribosome hibernation factors such as Lso1 and Lso2 (77, 78) may similarly have functions in protein translation or ribosome production in exponentially growing cells.

Despite the low number of mature ribosomes, *stm1Δ* cells do not show a significantly reduced growth rate, suggesting that the remaining ribosomes may be involved in translation (79). Consistently, the ribosome polysome fractions are comparable in wild-type and *stm1Δ* cells (**Fig 4G and H**). Additionally, it is possible that Stm1 shortens the time for re-engagement of individual ribosomes in the translation cycle, thus the fraction of translating ribosomes remains unchanged even though total levels of ribosomes are decreased. Alternatively, translational activity of individual ribosomes may be increased in the absence of Stm1. Indeed, Stm1 may repress translation by preventing association of elongation factor eEF3 with ribosomes (75). Further work measuring translational output and the activity of single ribosomes will be required to distinguish these possibilities.

### SERBP1 may similarly regulate ribosome homeostasis in mammalian cells

Mammalian cells express the Stm1 homolog SERPINE mRNA binding protein 1 (SERBP1), which like Stm1 binds 80S ribosomes in the cytosol inhibiting subunit splitting and translation activity (32, 80) and accumulates in the nucleolus in response to multiple stress conditions (81). The high sequence bias towards arginine, glycine, and glutamine amino acids and the presence of RGG/RGX motifs suggest that SERBP1 binds RNA and partitions into membrane-less organelles such as the nucleolus and nuclear speckles (82). Indeed, SERBP1 interacts with RiBi factors and may function in ribosome biogenesis and rRNA modification (83). Moreover, mammalian cells express HABP4, which shares many sequence motifs with SERBP1 and may thus perform redundant functions (84). Interestingly, SERBP1 emerged as oncogene for glioblastoma (GBM) development and increased expression in GBMs is associated with poor prognosis (85). SERBP1 may join the growing list of proteins linked to ribosome imbalances, ribosomopathies, disorders in which genetic abnormalities cause impaired ribosome biogenesis and function. Given the conserved ribosome functions of Stm1 and SERBP1, our findings may be relevant for future studies in higher and more complex eukaryotes to unravel mechanisms of ribosome homeostasis and ribosome crosstalk across multiple cellular compartments.

## Author contributions

Conceptualization: E.B. and M. Peter; formal analysis: E.B; investigation: E.B., M.B., F.U., M.W., J.T., S.Z., B.A., M.O.-O., R.D., J.H., P.K.; writing – original draft: E.B. and M. Peter; writing-review and editing: E.B., M. Peter with input from all authors; visualization: E.B., M. Peter; supervision: V.G.P., M. Peter and M. Pilhofer; funding acquisition: V.G.P, M. Pilhofer, M. Peter.

## Acknowledgements

We thank Tom Dever for eIF2-alpha antibody and David Shore for his critical input; we are grateful to Jingwei Xu for establishing the pipeline to estimate ribosome concentration from cryo-tomographic data, Ludovic Gillet for mass spectrometry support, Prashant Rawat and the Peter lab for input and helpful discussions. We are grateful to ScopeM for instrument access, FCGZ and Lennart Opitz for sequencing and data processing services, to Alicia Smith for critical editing of the manuscript. Work in the Peter laboratory was supported by the Swiss National Science Foundation (SNSF) and ETH Zürich. The Pilhofer Lab was supported by the NOMIS foundation.

## Data and code availability

All the MS raw and analyzed data files, as well as scripts were deposited on Proteomics IDEntifications Database (PRIDE): PXD046775, PXD046821 for APMS of Stm1ΔIDR and Stm1ΔN, respectively; PXD046821 for targeted analysis of CURI complex components with Stm1ΔIDR, and PXD046820 for the pulsed-SILAC dataset). RNA Pol II Ser5 ChIP-Seq FASTQ raw files were deposited on the European nucleotide Archive (ENA): PRJEB72705.

## Supplementary figure legends

**Supplementary Figure 1.**
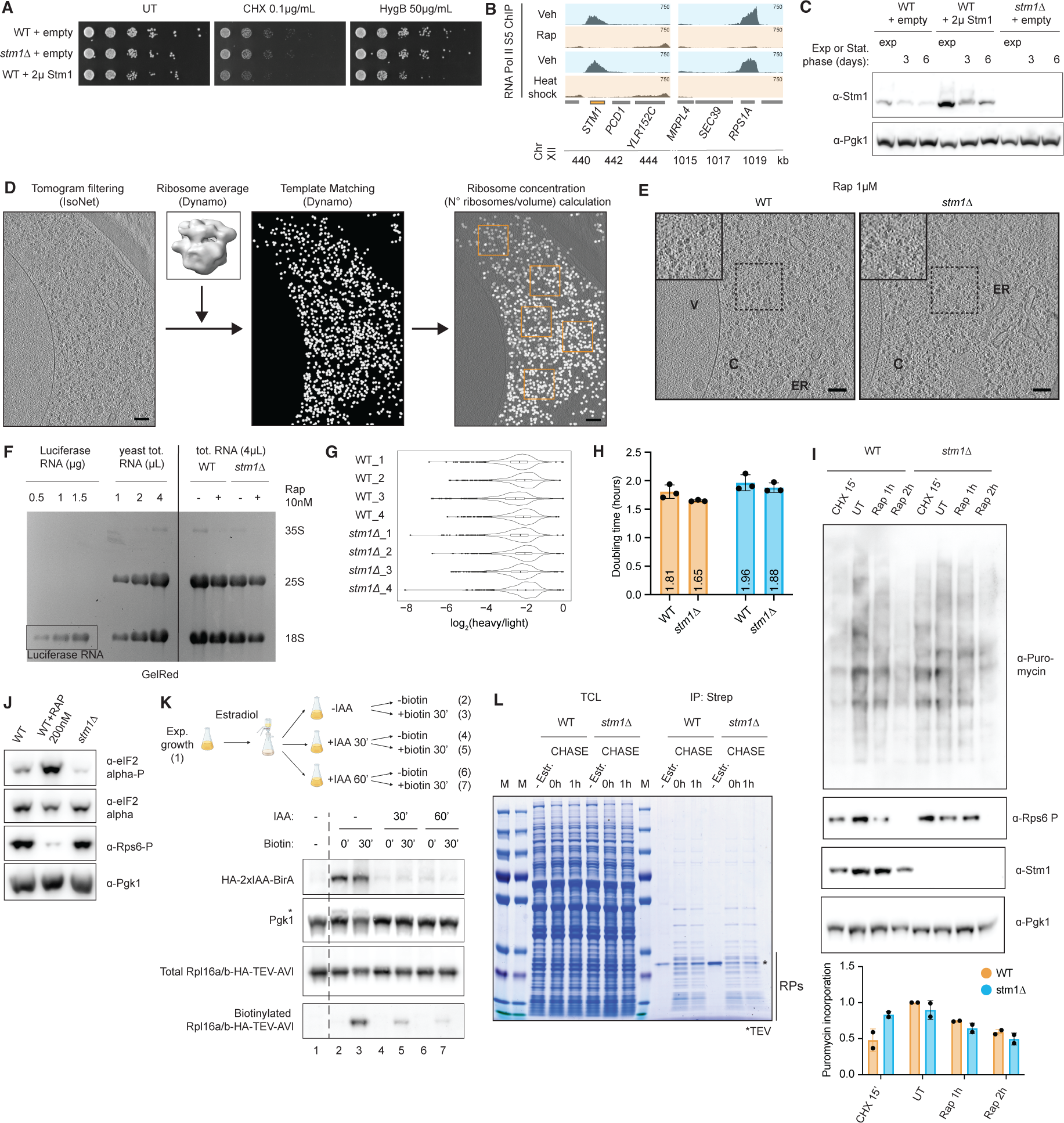
**A)** Serial dilution spotting assay monitoring growth of WT and *stm1Δ* cells carrying an empty control plasmid (+ empty) or a 2µ plasmid overexpressing Stm1 (+2µ Stm1) on synthetic auxotrophic SD-URA media. Cells were treated or not (UT) with cycloheximide (CHX 0.1µg/mL) or hygromycin B (HygB 50µg/mL) and incubated for 3 days. A representative image from N=3 independent experiments is shown. **B)** Distribution of Chip-Seq reads for RNA Pol II Ser 5 occupancy. The abundance of the *STM1* gene was compared to other Chromosome XII genes (PCD1, YLR152C, MRPL4, SEC39), before (Veh), after 20 minutes of rapamycin treatment (200nM) (Rap), and after heat shock for 5 min at 40°C. Data are re-analyzed from the Chip-Seq RNA Pol II dataset (GSE125226) (9). **C)** Western blot of total cell lysates prepared from WT or *stm1Δ* cells carrying an empty control plasmid (+ empty) or a 2µ plasmid overexpressing Stm1 (+2µ Stm1). Samples were collected from exponential growth phase (exp) or after 3 or 6 days of stationary phase (Stat. phase (days)). Blots were probed with antibodies against Stm1 and Pgk1 to control equal loading. N=2 independent replicates. **D)** Schematic representation of the iterative workflow to calculate riboosome concentration in cryo-tomograms. Briefly, tomograms were Contrast Transfer Function (CTF) deconvolved using Isonet (87) and used for template matching analysis with Dynamo (74). The average structure of a cytoplasmic ribosome was used as template reference. To calculate the ribosome concentration, five boxes of equal volume were randomly placed in the cytoplasm of each tomogram and used to count ribosomes. The number of coordinate points within each box was then used to calculate the molar concentration of cytosolic ribosomes. For more details, please refer to Methods. **E)** Representative cryo-tomograms of rapamycin (Rap 1µM for 2 hours) treated WT (left panel) or *stm1Δ* (right panel) cells, which were subjected to plunge freezing and focused ion beam (FIB)-milling, prior to cryo-electron tomography. The region of interest (dashed box) is shown magnified as an inset (solid black box) in the upper left corner. ER: endoplasmic reticulum, V: vacuole. Scale bars: 100 nm. Quantification of these images is shown in **Fig 1D.** **F)** GelRed-stained agarose gel analysis of total RNA samples (tot. RNA) from exponentially growing WT and *stm1Δ* cells. The cell number was normalized by OD_600_, and equivalent sample volume was loaded (4 µL). Cells were untreated (-) or treated (+) with rapamycin (Rap 10nM) for 2 hours. As standards, increasing amounts of Luciferase mRNA (Luciferase RNA (µg)), and increasing volume of total yeast RNA (yeast tot. RNA (µL)) were loaded on the agarose gel (left). The positions of the 35S, 25S, 18S and Luciferase RNA are marked. Quantification of N=3 independent experiments is shown in **Fig 1E.** **G)** Violin plot showing the ratio of proteins incorporating heavy and light lysine residues in WT and *stm1Δ* cells (N=4). Linked to **Fig 1F**. **H)** Bar plot showing the doubling time extracted from growth curves of WT and *stm1Δ* cells grown in SD media containing light lysine (orange) or grown in light lysine and then switched to SD media containing heavy lysine (blue). Data values are shown as dot as well as the mean and the standard deviation. N=3 independent replicates. **I)** Western blot of total cell lysates prepared from WT and *stm1Δ* cells treated with cycloheximide 200µg/ml for 15’ (CHX 15’), rapamycin 10nM for 1 hour or 2 hours (Rap 1h, Rap 2h) or left untreated (UT) prior to exposure to puromycin for the last 15’ of treatment. Blots were probed with antibodies to puromycin, phospho-Ser232/233 Rps6, Stm1 and Pgk1 (loading control). Bar plot quantification of puromycin incorporation normalized to the Pgk1 loading control is shown in the lower panel. Data values are shown as dots as well as the mean and standard deviation. Representative blot of N=2 independent replicates. **J)** Western blot of total cell lysates prepared from WT and *stm1Δ* cells treated or not with 200nM rapamycin for 2 hours (WT + Rap 200nM). Blots were probed with antibodies against eIF2alpha (Sui2), phospho-eIF2alpha (Sui2 Ser51), phospho-Rps6 (Ser232/233), or Pgk1 to control equal loading. N=3 independent replicates. Linked to **Fig 4B.** **K)** Schematic representation of ribosome biotinylation assay (upper panel). Cells were grown to exponential phase in low-biotin media with and then supplemented with 5 nM estradiol for 2 hours to induce BirA expression. Cells were filter-washed and split in three flasks. One flask was treated with auxin 1mM for 30’ (+IAA 30’) and another one for 60’ (+IAA 60’) to promote BirA degradation, while the last one was left untreated (-IAA). Each flask was then supplemented with biotin for 30’ (biotin 30’) to induce biotinylation or left untreated for control. Protein samples were collected before estradiol induction (lane 1) and after each auxin and/or biotin treatment (lanes 2-7). Western blot of total cell lysates prepared from cells expressing BirA tagged at its N-terminus with HA-2xIAA, and Rpl16a/b C-terminally tagged at its endogenous locus with HA-TEV-AVI (lower panel). Blots were probed with HA-antibodies to detect HA-2xIAA-BirA and total Rpl16a/b-HA-TEV-AVI. HRP-conjugated streptavidin was used to visualize biotinylated Rpl16a/b-HA-TEV-AVI (Biotinylated Rpl16a/b-HA-TEV-AVI), and Pgk1 controls equal loading. Representative blot of N=3 independent replicates. **L)** Coomassie stain of protein samples used in Figure 1I. M: protein ladder. *TEV: TEV protease treatment, RPs: ribosomal proteins. Representative gel from N=3 independent replicates.

**Supplementary Figure 2.**
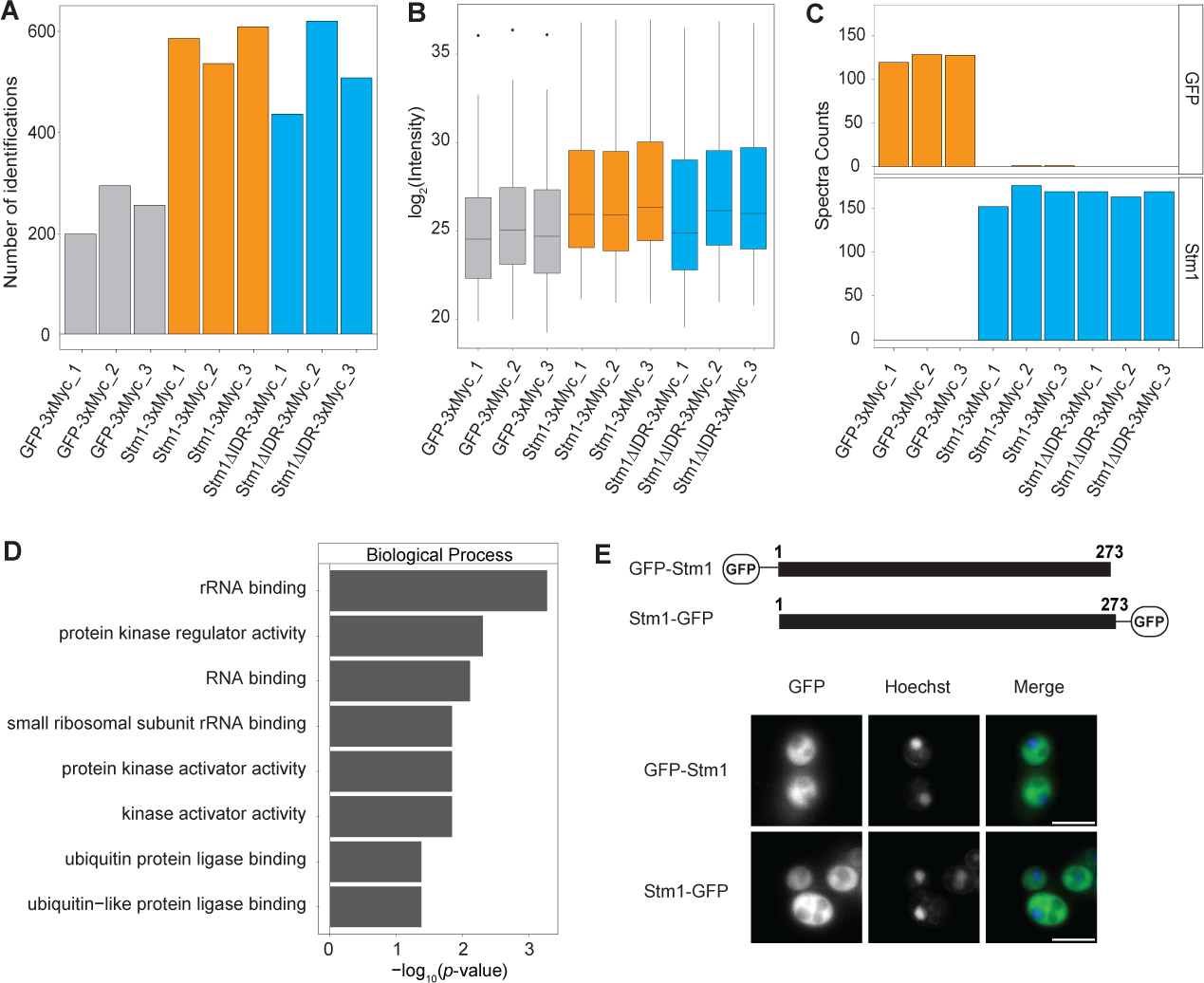
**A)** Bar plot displaying the number of protein identifications across all replicates for AP-MS analysis of GFP-3xMyc, Stm1-3xMyc and Stm1ΔIDR*-*3xMyc as in **Fig 2A**. N=3 independent replicates. **B)** Boxplot showing the log_2_ intensity across all replicates for AP-MS analysis of GFP-3xMyc, Stm1-3xMyc and Stm1ΔIDR-3xMyc as in **Fig 2A**. N=3 independent replicates. **C)** Barplot displaying the spectra counts across all replicates for AP-MS analysis of GFP and Stm1 in the affinity purification and MS analysis of GFP-3xMyc, Stm1-3xMyc and Stm1ΔIDR-3xMyc as in **Fig 2A**. N=3 independent replicates. **D)** Gene ontology for Biological Process calculated for Stm1-3xMyc protein interactors identified in **Fig 2A** (defined as log_2_FC> 2 and *p*-value< 0.05). The significance of the enrichment is reported as the -log_10_ of *p*-value. **E)** Representative wide-field light microscopy images of live cells stably expressing GFP-tagged Stm1, in which the GFP is inserted at either the N- (GFP-Stm1) or C-terminus (Stm1-GFP). Nuclei were stained with Hoechst. Scale bar: 5µm. N=2 independent replicates.

**Supplementary Figure 3.**
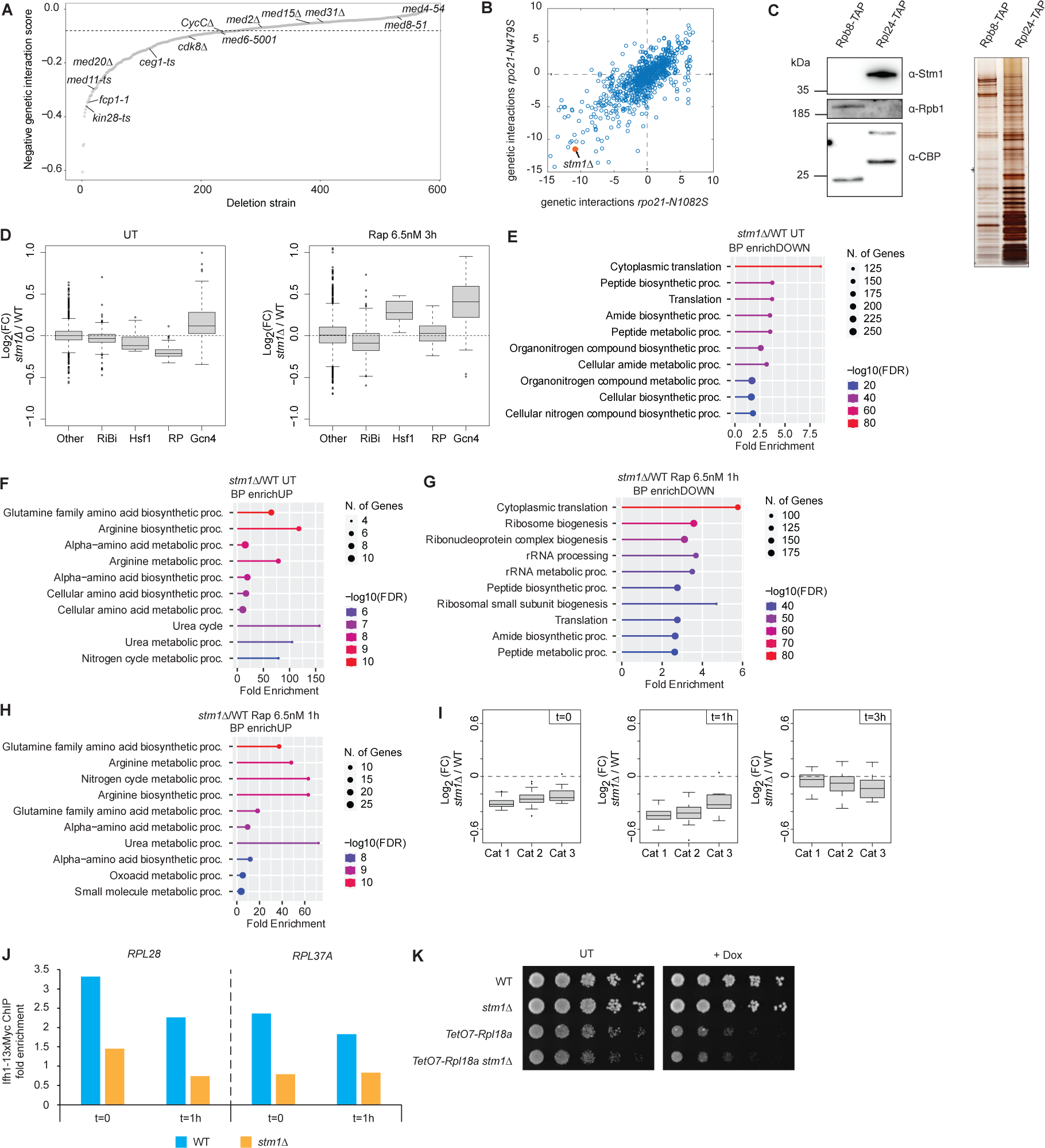
**A)** Distribution of the negative genetic interaction score of *stm1D* cells analyzed from the DRYGIN repository (43). The DRYGIN default cut off for intermediate genetic interaction strength is highlighted by the dashed line at y= 0.08. Components of the RNA Pol II machinery are labelled. **B)** Correlation plot of *rpo21-N479S* and *rpo21-N1082S* genetic interaction maps analyzed from (44). *STM1* deletion (*stm1Δ*) is highlighted in orange. **C)** Tandem affinity purification of the RNA Pol II subunits Rpb8-TAP and 60S ribosomal protein Rpl24-TAP. Western blots (left panel) and a silver-stained gel (right panel) are shown. Blots were probed with antibodies against CBP (Rpb8-TAP and Rpl24-TAP), Stm1 and Rpb1/Rpo21. N=2 independent replicates. **D)** Box plots showing RNA Pol II Ser 5 occupancy. The differential analysis of *stm1Δ* and WT cells is shown for ribosomal protein genes (RP (n=138)), ribosome biogenesis genes (RiBi (n=566)), Gcn4 targets genes (Gcn4 (n=108)), heat shock genes (Hsf1 (n=18)) and others (n=4311). Data are displayed as the log_2_ normalized fold change of RNA Pol II S5 occupancy in *stm1Δ* versus WT cells, either using untreated conditions (UT, left panel) or after 3 hours exposure to low rapamycin concentration (Rap 6.5nM, right panel). The signal was quantified for each RNA Pol II transcribed gene between the transcription start (TSS) and termination site (TTS). The gene categories have been defined according to scientific literature: (56) for RPG, RiBi, (86) for Hsf1, (47) for Gcn4 targets. An independent replicate is shown in **Fig 3B**. **E)** and **F**) Gene ontology enrichment for biological process calculated for genes with downregulated (**E**) and upregulated (**F**) RNA Pol II occupancy in *stm1Δ* versus WT cells in untreated conditions defined as log_2_FC< -0.5 and false discovery rate FDR< 0.001. The x-axis shows the enrichment compared to the whole yeast genome, the size of the circle represents the number of genes, and the color is indicative for the significance of the enrichment expressed as -log_10_(FDR). The analysis was performed with ShinyGO (version 0.76) (88) from data linked to **Fig 2B and Sup 3D.** **G) and H)** Gene ontology enrichment for Biological process calculated for genes with downregulated (G) upregulated (H) RNA Pol II occupancy in *stm1Δ* versus WT cells in rapamycin conditions (Rap 6.5nM 1h) defined as log_2_FC< -0.5 and false discovery rate FDR< 0.001. The x-axis shows the enrichment compared to the whole yeast genome, the size of the circle represents the number of genes, and the color is indicative for the significance of the enrichment expressed as -log_10_(FDR). The analysis was performed with ShinyGO (version 0.76) (88) from data linked to Fig 2B and Sup 3D. **I)** Box plots showin RNA Pol II Ser 5 occupancy change in *stm1Δ* versus WT cells for ribosomal protein genes (RP (n=138)) categories number 1 (Cat 1), number 2 (Cat 2) andumber 3 (Cat 3) (56). Data from biological replicate N=2 are displayed as the log_2_ normalized fold change of RNA Pol II occupancy in *stm1Δ* versus WT cells after no treatment (t=0, left panel) or treatment with 6.5 nM rapamycin for 1h (t=1, central panel) or 3h (t=3, right panel). An independent replicate is shown in **Fig 3D.** **J)** Bar plot showing Ifh1-13xMyc occupancy change analyzed by ChIP-qPCR in *stm1Δ* versus WT cells for the RPGs *RPL28* and *RPL37A* in untreated (t=0) and treatment with 6.5nM rapamycin for 1 h (t=1h) normalized on *ACT1*. An independent replicate is shown in **Fig 3E.** **K)** Serial dilution spotting assay to monitor growth on YPD media of WT and *stm1Δ* cells, and WT or *stm1Δ* cells where the endogenous promoter of *RPL18A* was replaced by a doxycycline repressible promoter (*TetO7-Rpl18a*). Cells were treated or not (UT) with 1µg/mL doxycycline (+ Dox) and imaged after 3 days. N=3 independent experiments.

**Supplementary Figure 4.**
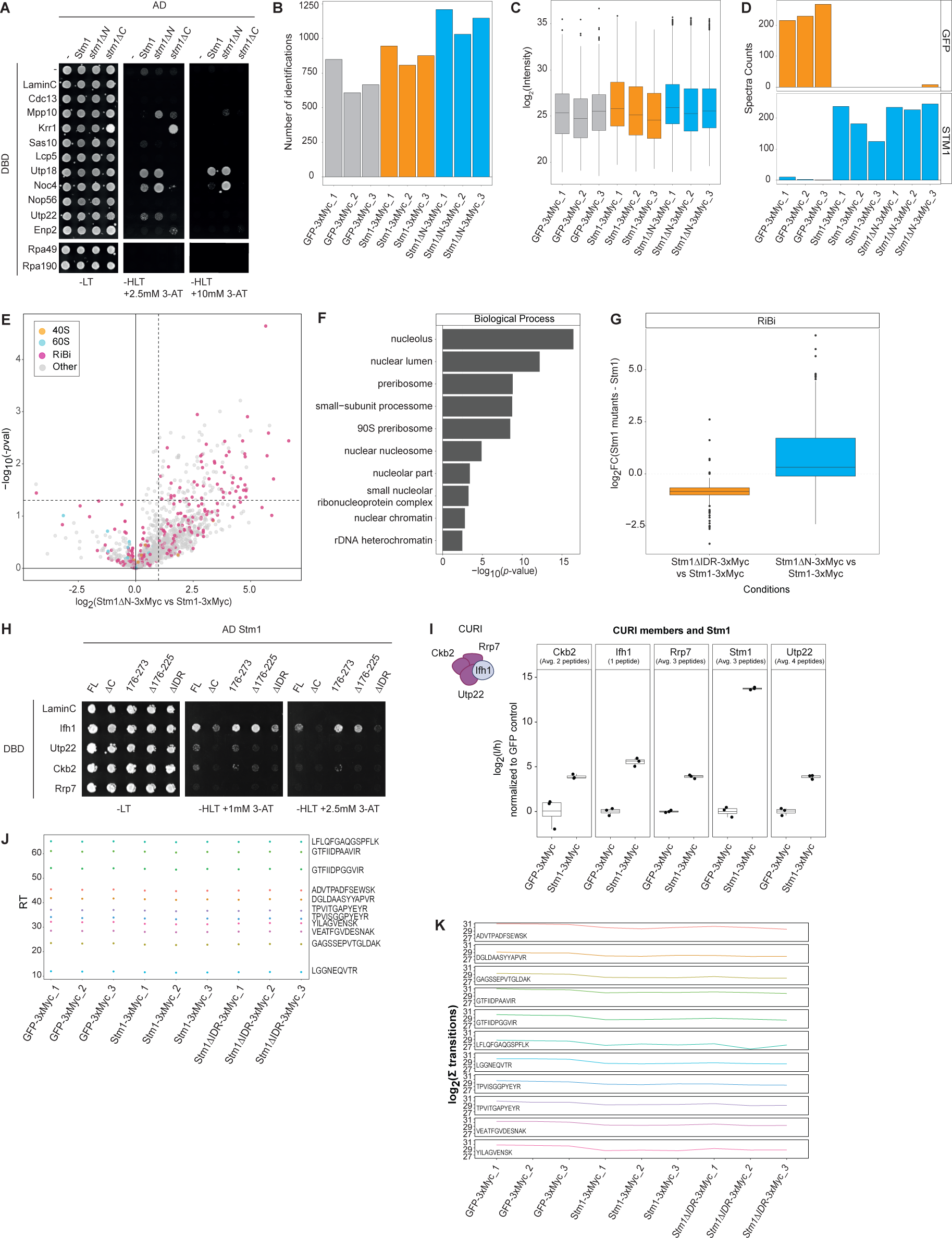
**A)** Two-hybrid analysis of full length Stm1 (Stm1) and N- (*stm1ΔN)* or C-terminally (*stm1Δ*C) truncated Stm1 fused to the GAL4 activation domain (AD) with components of the SSU fused to GAL4 DNA binding domain (DBD). For control, unfused AD or DBD domains (-), or DBD-fusions to Lamin C and Cdc13 were included. Colonies were grown on permissive auxotrophic SD -Leu, -Trp (-LT), or selective -His, -Leu, - Trp agar media, supplemented with 2.5mM or 10mM 3-amino-1,2,4-triazole (-HLT +3-AT). Representative picture of N=3 independent replicates. Extension of **Fig 2E.** **B)** Bar plot displaying the number of protein identifications across all conditions for AP-MS of GFP-3xMyc, Stm1-3xMyc and Stm1ΔN-3xMyc. Linked to **Sup 4E.** **C)** Box plot showing the distribution of protein intensities across all conditions for AP-MS of GFP-3xMyc, Stm1-3xMyc and Stm1ΔN-3xMyc. Linked to **Sup 4E**. **D)** Bar plot displaying the spectra counts across all conditions for AP-MS of GFP-3xMyc, Stm1-3xMyc and Stm1ΔN-3xMyc. Data values for each biological replicate are shown (N=3). Linked to **Sup 4E**. **E)** Volcano plot showing the log_2_ fold change enrichment of proteins co-purified with Stm1ΔN-3xMyc compared to Stm1-3xMyc. The protein abundance was normalized for Stm1 levels after AP-MS. The significance is calculated using a two-sided unpaired Student’s t-test. N=3 independent replicates. Components of the 40S (orange) and 60S (blue) ribosomal subunits and RiBi factors (purple) are highlighted. **F)** Gene ontology for Biological Process was calculated for the Stm1 interactors identified in **Sup 4E** defined as log_2_FC > 1 and *p-*value< 0.05. The significance of the enrichment is reported as the -log_10_ of *p*-value. **G)** Box plot of the log_2_ fold change enrichment of RiBi proteins identified in the AP-MS analysis of the Stm1 N- (Stm1ΔN-3xMyc) and C-terminal (Stm1ΔIDR-3xMyc) truncation mutants, relative to full-length Stm1-3xMyc. Linked to **Sup 4E and Fig 5A.** **H)** Two-hybrid analysis of full length Stm1 (FL), Stm1ΔC, Stm1ΔIDR, Stm1Δ176-225, and the Stm1 C-terminus (AA176-273) fused to GAL4 activation domain (AD) with components of the CURI complex fused to GAL4 DNA binding domain (DBD). The DBD-LaminC fusion was included for negative control. Cells were grown on permissive auxotrophic SD -Leu, -Trp (-LT), or selective -His, -Leu, -Trp agar media, supplemented with 1mM or 2.5mM 3-amino-1,2,4-triazole (-HLT +3-AT). Cells were imaged after 3 days of incubation. Representative image of N=3 independent replicates. **I)** Targeted PRM measurement of Ifh1 and CURI subunits co-purified with Stm1-3xMyc. Box plots showing the log_2_ fold change enrichment of Ckb2, Ifh1, Rrp7 and Utp22 after AP-MS analysis of Stm1-3xMyc relative to GFP-3xMyc controls (**Fig 2A**). Data values are shown as log_2_ ratio of light (endogenous peptide) to heavy (stable heavy isotope-labeled peptide, reference peptides) (l/h). Data values show protein abundance calculated from the average intensity of measured peptides (the intensity of all measured peptides is reported in **Sup 5A, Supplementary Table 5**). The number of peptides used for each protein is indicated below the protein name. Each dot represents an independent replicate (N=3), with the box plot boundaries indicating the quantiles Q1 (25%) and Q3 (75%) and the average value denoted by a line across the box (N=3). **J)** Quality control for targeted analysis reporting the retention time (RT) of iRT peptides (internal standards for targeted analysis). Linked to **Sup 4I**. **K)** Quality control for targeted analysis reporting intensity of iRT peptides (internal standards for targeted analysis). The intensity of the peptides is obtained from the sum of all peptide transitions. Linked to **Sup 4I.**

**Supplementary Figure 5.**
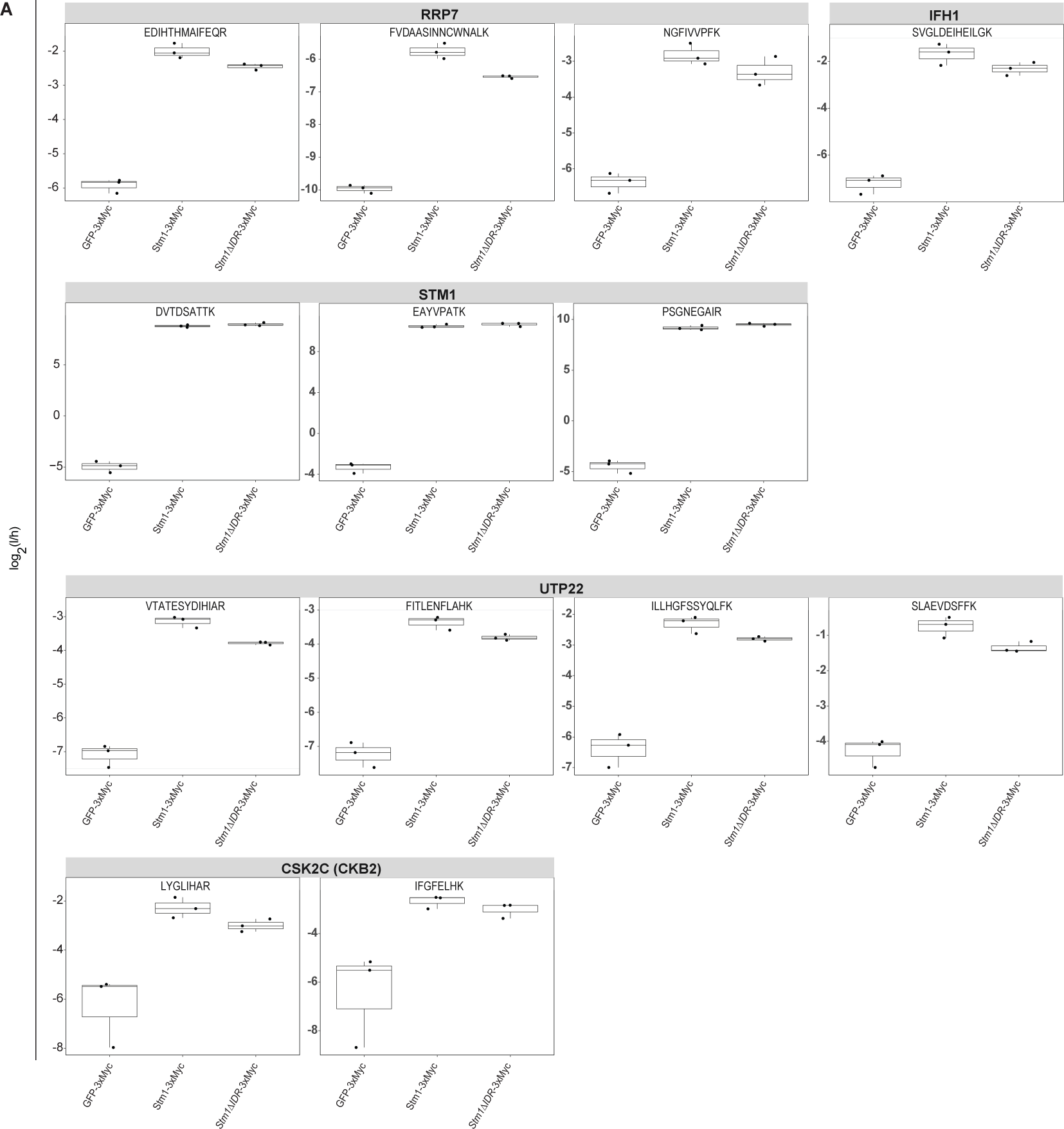
**A)** Targeted PRM measurement of Ifh1 and CURI complex members co-precipitating with Stm1-3xMyc. This panel reports the log_2_FC enrichment for all measured peptides in AP-MS measurements of Stm1-3xMyc compared to the GFP-3xMyc control (**Fig 5C, Sup 4I**). Data values are shown as log_2_ ratio of light (endogenous peptide) to heavy (stable heavy isotope-labeled peptide) (l/h). Each dot represents the results for a technical replicate (N=3), with the box plot boundaries indicating the quantiles Q1 (25%) and Q3 (75%) and the average value denoted by a line across the box (N=3). Linked to **Sup 4I.**

## Material and methods

### Strains and plasmids

All plasmids and strains used in this study are listed in **Supplementary Table 6**. Plasmids were generated using standard molecular biology methods and checked by sequencing. Yeast strains are derivatives of BY4741, BY4742, W303-1A and NMY32 backgrounds. Gene replacement and tagging at the endogenous locus were obtained by PCR-based homology recombination or by crossing and sporulation. Cells were grown at 30°C in yeast extract pentose dextrose (YPD) (1% yeast extract, 2% peptone and 2% dextrose) or in synthetic defined (SD) medium (0.17% Yeast nitrogen base without amino acids, 2% carbon source, 0.5% NH_4_ sulfate and 1x amino acids mix) and harvested in mid-log phase (OD_600_ = 0.6), unless differently stated.

### Spotting assay

Cells were inoculated overnight. The next day, cultures were diluted to OD_600_ = 0.1-0.15. After two generations cells were normalized to OD_600_ = 0.3. Five 1:5 serial dilutions were prepared in a 96-well round bottom plate (50µL cells suspension in 250µL total volume). Dilutions were spotted on YPD or SD agar plates containing rapamycin (LC Laboratories, #R-5000), cycloheximide (Sigma, #C7698), Hygromycin B (Corning, #30-240-CR) at the indicated concentrations. Pictures were taken after growth for 2-3 days at 30^°^C.

### Growth curves and doubling times

Cells were inoculated in SD without L-Lysine (SD - Lys) media but supplemented with light L-Lysine (25mg/L) and diluted after 8h into exponential phase. After 20 hours, OD_600_ was measured and normalized to 0.1. Cells to be grown in SD media containing heavy Lysine (^13^C_6_ ^15^N_2_ (Cambridge Isotope Laboratories #CNLM-291-H) (25mg/L) were filter washed using a 0.45µM pore size nitrocellulose membrane (Millipore), washed twice with SD + heavy Lysine and resuspended in SD + heavy Lysine. The same procedure was executed with SD + light Lysine for cells to be grown in light Lysine. The optical density was recorded over time using a plate reader (Spectrostar BMG Labtech). Measurements of 3 biological independent replicates are plotted as mean and standard deviation in GraphPad Prism (version 9.5.1). Doubling times were extracted from exponential growth phase, and calculated using the Prism in-built nonlinear regression equation for exponential growth:

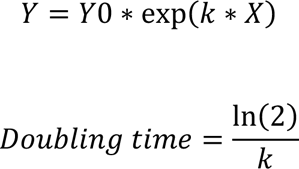

Y0 is the initial Y value, X the time expressed in hours and k the rate constant. Y is expressed as hours^-1^.

### Two-Hybrid assay

Full-length ORFs were amplified by PCR from genomic DNA and cloned into pLexA (encoding for the DNA binding domain) or pACT2.2 plasmids (encoding for the activation domain) (Dual System AG) using restriction enzymes, mutagenesis or Gibson assembly (89). The correct integration into the multiple cloning site was verified by sequencing (Microsynth AG) and SnapGene software (5.2.2). Plasmids were co-transformed into NMY32 yeast cells. Clones originating from single cells were selected and grown overnight in SD media with auxotrophic selection (-Leu, -Trp). Cells suspensions were then diluted into fresh media and allowed to reach exponential phase. A single dilution of cells (OD_600_ = 0.3) was spotted on SC -Leu, -Trp (-LT) or SC -His, -Leu, -Trp (-HLT) agar plates containing 3-Amino-1,2,4-triazole (Sigma, A8056) at the indicated dilutions. Pictures were taken after growth at 30^°^C for 2-4 days.

### β-Galactosidase assay

β-galactosidase expression was monitored using the PXG assay (90). Briefly, yeast cells were inoculated in selective medium (SD-Leu-Trp) and grown overnight at 30°C. Cultures were then diluted to OD_600_ = 0.2 and grown for approximately 5 hours to reach OD_600_ = 0.6-0.8. 1.4 OD of cells were pelleted by centrifugation at 2500 *g* for 3 min and resuspended in 130 μL X-gal reagent (0.1% (w/v) X-Gal (Sigma-Aldrich, #3117073001), 0.05% (v/v) β-mercaptoethanol and 0.1% (w/v) SDS in 1x PBS). 125 μL was transferred to a transparent flat bottom 96-well microplate and incubated at room temperature for several hours. Color changes were scanned at different time points using a flatbed scanner.

### Protein sample preparation and western blotting

Cells corresponding to 1 OD were resuspended in 1mL of ice-cold water, lysed on ice in 150 µL 2M NaOH with 0.7% (v/v) ϕ3-mercaptoethanol and precipitated with TCA solution (55% w/v). Protein samples were centrifugated at 16,000 *g* for 15 min at 4°C and resuspended in 33 µL of HU Buffer (120mM Tris pH 6.8, 5% glycerol, 8M urea, 0.1% bromophenol blue, 3% (v/v) ϕ3-mercaptoethanol freshly added) and boiled at 65°C for 15 minutes.

Protein samples were loaded on polyacrylamide Bolt TM 4-12%, Bis-Tris gradient gels (Invitrogen, #NP0321) and separated at 120-150V in 1X MOPS or MES buffer. Proteins were then transferred on polyvinylidene (PVDF) (Sigma #P2938) or nitrocellulose (Amersham, #10600002) membranes by electro-transfer. Protein transfer was confirmed using Ponceau Red staining (0.5% (w/v), Ponceau S dissolved in 1% (v/v) acetic acid). Membranes were blocked in 1x PBS-Tween20 (0.05%) containing 5% milk powder or 1x TBS-Tween (0.05%) with 5% BSA. The following primary antibodies were used: Pgk1 (1:3000, α-mouse, Life technologies, #459250), c-Myc (1:3000, α-mouse, Thermo Fisher Scientific, #MA1-980), CBP (1:2000, α-rabbit, Upstate, #07-482), GFP (1:3000, α-mouse, Roche Diagnostics, #11814460 001), phospho-S6 (Ser235/236 corresponding to yeast Rps6 Ser 232/233) (1:2000, α-rabbit, Cell Signalling #4858), eIF2α (1:1000, α-rabbit, generously provided by T. Dever), phospho-eIF2α (1:3000, α-rabbit, Enzo Life Sciences, #BML-SA405), puromycin (1:5000, α-mouse, clone 12D10, Millipore #MABE343), Rpb1 (1:3000, α-mouse, Biolegend, #664906), Rps5 and Rpl35 (both 1:3000, α-rabbit, kind gifts from V. Panse). Stm1 antibody was raised in rabbit against the protein C-terminus (Eurogentec) and used at 1:2000 dilution. Secondary horseradish peroxidase-coupled antibodies (1:3000, Pierce, α-mouse #1858413, α-rabbit #1858415) were applied prior to ECL-based chemiluminescence (Biorad #1705061 and ThermoFisher #34094) detected using a Fusion Fx instrument (Vilber Lourmat). Biotinylated Rpl16a/b was detected by HRP-conjugated Streptavidin (1:10’000, ThermoFisher #N100).

### Ribosome polysome analysis by sucrose gradients

Sucrose gradient analysis of ribosome polysomes was performed adapting experimental conditions from (24). Briefly, 200mL of exponentially growing cells were treated with CHX 100µg/mL (Sigma, #C7698) freshly dissolved in DMSO (Sigma, #D8418) and cooled on ice for 2 minutes. Cells were harvested by centrifugation in conical 500mL bottles (2 min 2000 *g*), washed once with 20mL ice cold 1X PBS containing CHX 100µg/mL and once with lysis buffer (20 mM Tris–HCl pH 7.5, 150 mM KCl, 5 mM MgCl_2_, 1% Triton X100, 1 mM DTT, 100 μg/ml CHX, 0.1 mM PMSF) and lysed in 1mL lysis buffer and 0.2g (400-600µM) glass beads by vortexing (Disruptor Genie, Scientific Industries) for 15min. After recovering the lysates in a fresh tube, they were clarified by centrifugation at 4°C for 10 minutes, top speed. RNA concentration at 260nm was measured and aliquots of 4 units of A_260_ were supplemented with 5% glycerol and flash-frozen in liquid nitrogen. Each aliquot was layered on a 10-50% sucrose gradient in 20 mM Tris–HCl pH 7.5, 150 mM KCl, 5 mM MgCl_2_, 1 mM DTT, 100 μg/ml CHX, 0.1 mM PMSF buffer and subjected to ultracentrifugation at 36,000 rpm at 4°C for 2h using a SW41 rotor (Beckman) and 14 x 89 mm polypropylene tubes (Beckman, #331372). Profiles were recorded at 260nm with a fractionator device (Biocomp). RNA absorbance A_260_ was recorded and plotted with Excel.

### Chromatin-immunoprecipitation, sequencing analysis and qPCR

Yeast cells were grown in YPD medium at 30°C overnight before diluting the cultures to OD_600_ = 0.1. The cells were grown to log-phase (OD_600_ = 0.3–0.4) and treated with 6.5 nM rapamycin final concentration (10 μM of stock solution resuspended in 90% ethanol, 10% Tween-20) for 60 min before crosslinking. ChIP experiments were performed with exponential phase cells harvested around OD_600_ β 0.6-0.8. Yeast cultures of 100 mL in complete medium were collected and then crosslinked with 1% (v/v) formaldehyde for 15 min and quenched by adding 125 mM glycine for 5 min at room temperature. Cells were then washed with ice-cold HBS (50 mM HEPES-Na pH7.5, 140 mM NaCl) and resuspended in 0.6 mL of ChIP lysis buffer (50 mM HEPES-Na pH7.5, 140 mM NaCl, 1 mM EDTA, 1% NP-40, 0.1% sodium deoxycholate) supplemented with 1 mM PMSF and 1x protease inhibitor cocktail (Roche). The cells were broken using Zirconia/Silica beads (BioSpec) and lysates centrifuged at 13,000 rpm for 30 min at 4°C. Pellets were resuspended in 300 μL ChIP lysis buffer containing 1 mM PMSF and sonicated for 15 min (30s ON - 60s OFF) in a Bioruptor (Diagenode). The lysates were then centrifuged at 7000 rpm for 15 min at 4°C and soluble fractions were transferred to new tubes. A defined amount (5% of total chromatin) of crosslinked and sonicated chromatin from *S. pombe* was added to *S. cerevisiae* chromatin prior to the immunoprecipitation step. ChIP was performed by using RNAPII (Abcam, #ab5131) and Myc (Abcam, #ab32) antibodies and incubated overnight at 4°C with rotation. Magnetic beads coupled to IgG against rabbit or mouse (Invitrogen, Dynabeads™ M-280 Sheep Anti-Rabbit or Anti-Mouse IgG) were washed three times with PBS (137 mM NaCl, 2.7 mM KCl, 10 mM Na_2_HPO_4,_ 1.8 mM KH_2_PO_4_) containing 0.5% BSA and added to the lysates (30 μL of beads/300 μL of cell lysate). The samples were incubated for 2 hr at 4°C with rotation. The beads were washed twice with AT1 buffer (50 mM HEPES-Na pH7.5, 140 mM NaCl, 1 mM EDTA, 0.03% SDS), once with AT2 buffer (50 mM HEPES-Na pH7.5, 1 M NaCl, 1 mM EDTA), once with AT3 buffer (20 mM Tris-Cl pH7.5, 250 mM LiCl, 1 mM EDTA, 0.5% NP-40, 0.5% sodium deoxycholate) and twice with TE buffer (10 mM Tris, 1 mM EDTA). Chromatin was eluted from the beads by resuspension in TE containing 1% SDS and incubation at 65°C for 10 min. The eluate was transferred to new eppendorf tubes and incubated overnight at 65°C to reverse the crosslinks. The DNAs were purified using the MinElute PCR Purification Kit (Qiagen).

DNA libraries were prepared using NEBnext Ultra according to manufacturer’s specifications. The libraries were sequenced with Illumina’s NovaSeq6000 at FGCZ (Zürich) in single read mode 100nt. The data analysis was performed as follows: the raw reads were first cleaned by removing adapter sequences, trimming low quality ends, and filtering reads with low quality (phred quality <20) using Fastp (Version 0.20) (91). Read alignment was done with Bowtie2 (v2.4.1) (92). As reference we used the Ensembl genome build R64 with the gene annotations downloaded on 2019-12-03 from Ensembl (release 98). Count values were computed with the featureCounts-function from the R package Rsubread (v2.4) (93). The options for featureCounts were: - min mapping quality 10 - min feature overlap 10bp - count multi-mapping reads - count only primary alignments - count reads also if they overlap multiple genes. Differentially expressed (DE) regions were identified using the R package edgeR (v3.30) (94) applying a generalised linear model (glm) regression, a quasi-likelihood (QL) differential expression test and the trimmed means of M-values (TMM) normalisation. The entire dataset is available in the ENA repository (PRJEB72705).

Gene categories of GCN4 targets, ribosomal protein genes category 1 to 3, RiBi factors and Hsf1 targets were defined according to previous studies (47, 56, 86). FASTQ files were mapped to the sacCer3 and the 972-h pombe genome assemblies using the Galaxy server (https://usegalaxy.org). For each sample, reads mapped uniquely to the sacCer3 genome were used for further analysis, while reads that mapped to both genomes were discarded. The signal was quantified for each gene between the transcription start site (TSS) and transcription termination site (TTS) for RNAPII. The gene list is reported in **Supplementary table 3**. For Ifh1-qPCR, after cross-linking, quantification of immunoprecipitated DNA was performed as described in (95) by real-time qPCR (Roche LightCycler 480). Ifh1 occupancy at the *RPL28* and *RPL37A* promoters was calculated as a fold enrichment relative to *ACT1* controls in *stm1Δ* and WT strains. Primer sequences for ChIP-qPCR assay are available upon request.

### Protein immunoprecipitation

For Streptavidin pull-down of biotinylated protein, a volume of cells corresponding to 80OD e.g. 100mL of OD_600_ = 0.8 cells in mid-log phase, were treated with 100µg/mL CHX and cooled on ice for 5 minutes. Cells were harvested by centrifugation in conical bottles (3500 *g* 4°C), washed and disrupted with glass beads in 800µL lysis buffer (50mM Tris-HCl pH7.5, 100mM NaCl, 1mM EDTA pH 8.0, 20mM MgCl_2_, 100µg/mL CHX, 1mM PMSF, cOmplete^TM^ EDTA free protease inhibitor (Roche, #11836170001), NP40 0.1%). Lysates were spun at 20’000 *g* for 20min at 4°C and the recovered supernatant was transferred to a new tube. After measuring protein concentration, lysates were normalized and 20µL input sample was taken. Lysates were incubated for 1 hour with 50µL of slurry Streptavidin-conjugated magnetic beads (Invitrogen, #65605D) rotating at 4°C. Beads were recovered with a magnet and washed 5 times in lysis buffer with 100µg/mL CHX, 1mM PMSF (without NP40 and cOmplete^TM^ EDTA free protease inhibitor). Beads were then resuspended in 40µL of lysis buffer with CHX and PMSF (without NP40 and cOmplete^TM^ EDTA free protease inhibitor) before adding TEV protease. Samples were rotated overnight at 4°C or at 16°C for 1h. Beads were recovered with the magnet, the supernatant transferred to a fresh tube and mixed with 40µL urea loading buffer (120mM Tris pH 6.8, 5% glycerol, 8M urea, 0.1% bromophenol blue, 3% (v/v) β-mercaptoethanol). Samples were heated at 65°C for 15 minutes before Western blotting analysis or SDS-PAGE and Coomassie staining.

Extracts for immunoprecipitation were prepared from 100mL of OD_600_ = 0.6-0.8 cells expressing Myc-tagged proteins, which were treated with 100µg/mL CHX and cooled on ice for 5 minutes. Cells were harvested by centrifugation in conical bottles (3500 *g*, 4°C), washed and disrupted with glass beads in lysis buffer (50mM Tris-HCl pH7.5, 100mM NaCl, 1mM EDTA pH 8.0, 20mM MgCl_2_, 100µg/mL CHX, 1mM PMSF, cOmplete^TM^ EDTA free) in a 1.5mL tube. After addition of NP40 0.5% (Sigma, #127087-87-0), lysates were spun on a tabletop centrifuge (20’000 *g* for 15min at 4°C) and cleared with GST agarose beads for 20 minutes. Anti-c-Myc magnetic beads (Thermo Fisher Scientific, #88843) were washed in lysis buffer and added to the lysates for 1h 30’ at 4°C with rotation. Beads were washed five times with lysis buffer supplemented with CHX and PMSF (without NP40), boiled in 50µL 1X SDS loading buffer and analyzed by Western blotting.

### Proteomics sample preparation

After Myc-affinity purification, samples were digested in-gel (AP-MS comparison Stm1-3xMyc and Stm1ΔN-3xMyc) or on-beads (AP-MS comparison Stm1-3xMyc and Stm1ΔIDR-3xMyc). For in-gel digestion, proteins were separated/concentrated by SDS-PAGE, reduced (6.5mM DTT, 1 hour at 60°C), alkylated (54mM Iodoacetamide, 30min, RT) and digested with 1µg of Trypsin (Promega) (96). Before MS injection, peptides were subjected to C18 cleanup (C18 7-70µg, The Nest Group) following manufacture instructions, dried and re-suspend in 2% acetonitrile 0.1% formic acid.

For on-bead digestion, proteins were digested after purification using Filter Assisted Spin Proteolysis (FASP) (97) as previously described (98). Briefly, proteins were loaded in 10 kDa molecular weight cutoff spin column (Vivacon 500, Sartorious) and centrifuged at 9000 *g* for 15min until dryness. Samples were dissolved in urea 8M, reduced (5mM) and alkylated (10mM), washed with ammonium bicarbonate (0.1M) and proteolyzed (0.4µg of LysC (Wako) for 3 hours and 1.2µg of Trypsin (Promega) overnight). Peptides were collected in a tube by centrifugation (8000 *g*, 20min) and the spin column was additionally washed with MS water. Proteolysis reaction was quenched with 5% formic acid. Before MS injection, peptides were subjected to C18 cleanup (C18 7-70µg, The Nest Group) following manufacture instructions, dried and re-suspend in 2% acetonitrile 0.1% formic acid. For targeted analysis, isotope-labeled heavy peptides corresponding to the selected proteotypic peptides, containing either heavy lysine (13C(6) 15N(2)) or arginine (13C(6) 15N(4)) were spiked with RT peptides (Biognosys) to each sample before LC–MS/MS analysis. The inclusion list with analyzed target peptides is reported in **Supplementary Table 5**.

## Mass spectrometry data acquisition

### Untargeted differential interactome analysis of WT Stm1 and the ΔN terminal truncation

LC-MS/MS analysis was performed on an Orbitrap QExactive+ mass spectrometer (Thermo Fisher) coupled to an EASY-nLC-1000 liquid chromatography system (Thermo Fisher). Peptides were separated using a reverse phase column (75 µm ID x 400 mm New Objective, in-house packed with ReproSil Gold 120 C18, 1.9 µm, Dr. Maisch GmbH) across a gradient from 5 to 30% in 90 minutes (buffer A: 0.1% (v/v) formic acid; buffer B: 0.1% (v/v) formic acid, 95% (v/v) acetonitrile). The DDA data acquisition mode was set to perform one MS1 scan followed by a maximum of 20 scans for the top 20 most intense peptides with MS1 scans (R=70’000 at 400 m/z, AGC=3e6 and maximum IT=64 ms), HCD fragmentation (NCE=30%), isolation windows (1.4 m/z) and MS2 scans (R=17’500 at 400 m/z, AGC=1e5 and maximum IT=55ms). A dynamic exclusion of 30s was applied and charge states lower than two and higher than seven were rejected for the isolation.

### Untargeted differential interactome analysis of WT Stm1 and the Stm1ΔIDR mutant

LC-MS/MS analysis was performed on an Orbitrap Exploris480 mass spectrometer (Thermo Fisher) coupled to an Vanquish Neo liquid chromatography system (Thermo Fisher). Peptides were separated using a reverse phase column (75 µm ID x 400 mm New Objective, in-house packed with ReproSil Gold 120 C18, 1.9 µm, Dr. Maisch GmbH) across a gradient from 7 to 35% in 120 minutes (buffer A: 0.1% (v/v) formic acid; buffer B: 0.1% (v/v) formic acid, 80% (v/v) acetonitrile). The DDA data acquisition mode was set to perform one MS1 scan followed by a maximum of 20 scans for the top 20 most intense peptides with MS1 scans (R=60’000 at 400 m/z, normalized AGC (%)=100 and max. IT set to auto), HCD fragmentation (NCE=28%), isolation windows (1.4 m/z) and MS2 scans (R=15’000 at 400 m/z, normalized AGC (%)=200 and max. IT set to auto). A dynamic exclusion of 25s was applied and charge states lower than two and higher than seven were rejected for the isolation.

### Targeted differential interactome analysis of Stm1 WT and the Stm1ΔIDR mutant

LC-MS/MS analysis was performed on an Orbitrap Exploris480 mass spectrometer (Thermo Fisher) coupled to an Vanquish Neo liquid chromatography system (Thermo Fisher). Peptides were separated using a reverse phase column (75 µm ID x 400 mm New Objective, in-house packed with ReproSil Gold 120 C18, 1.9 µm, Dr. Maisch GmbH) across a gradient from 7 to 35% in 60 minutes (buffer A: 0.1% (v/v) formic acid; buffer B: 0.1% (v/v) formic acid, 80% (v/v) acetonitrile). A targeted scheduled method was set up with one MS1 scan (R=120’000 at 400 m/z, normalized AGC (%)=100 and max. IT set to auto) and a cycle time of 3s to monitor 53 peptides (variable normalized AGC target and max IT) using an isolation window of 1.5m/z and normalized collision energy 27%.

### Mass spectrometry data analysis Untargeted analysis

For untargeted analysis, acquired spectra were searched using the MaxQuant software package version 1.5.2.8 embedded with the Andromeda search engine (99) against the *Saccharomyces cerevisiae* reference dataset (http://www.uniprot.org/, downloaded on 14.05.2020, 6’052 proteins) extended with reverse decoy sequences. The search parameters were set to include only full tryptic peptides, maximum two missed cleavage, carbamidomethyl as static peptide modification, oxidation (M) and deamidation (N-ter) as variable modification and “match between runs” option. The MS and MS/MS mass tolerance was set to 10 ppm, and a False discovery rate of < 1% was used protein identification. Protein abundance was determined from the intensity of top two peptides. Intensity values were median normalized and imputed using random sampling from a normal distribution generated 1% less intense values. ANOVA statistical analysis was employed to compare multiple condition. The entire dataset, including raw data, generated Tables and scripts used for the data analysis are available in the PRIDE repository (PXD046775 and PXD046821 for Stm1 delta IDR and Stm1 delta N terminal, respectively).

### Targeted analysis

For targeted analysis, peptides were analyzed manually using Skyline daily (100), and correct identification with six fragment ions per peptide was assigned based on the coelution of light and heavy peptide and matching peak shape for precursor and product ions from light and heavy peptides. Peptide abundance was analyzed by summing the integrated areas of three fragment ions per peptide. Significance of change in intensity was estimated with *p*-values using two-sided, not paired t-tests. The entire dataset, including raw data, generated tables and scripts for the data analysis are available in the PRIDE repository (PXD046821).

### Proteomics sample preparation for pulsed-SILAC experiment

Yeast cultures were grown at 30°C in SD with 25mg/mL light L-lysine in exponential phase for 24h. At OD_600_ = 0.4, cells were pulsed with heavy media. Cells were vacuum filtered on 0.45µm nitrocellulose membrane, washed with warm SD media containing heavy ^13^C_6_ ^15^N_2_ L-lysine 25mg/mL (Cambridge Isotope Laboratories #CNLM-291-H), and resuspended in warm SD heavy L-lysine. After 2h of pulse labelling, samples equivalent to 50mL OD_600_ = 0.8 were harvested by vacuum filtering on nitrocellulose membrane, washed with distilled water, scraped and snap frozen in liquid nitrogen. The samples where then processed for protein extraction and mass spectrometry analysis. Briefly, samples were resuspended in 100µL lysis buffer (8M urea, 100 mM Ammonium Bicarbonate, 5 mM EDTA, pH 8) and bead beaten with 200µL acid-washed glass beads. The samples were spun for 5 min with 500 *g* at 4°C and the supernatant transferred to a new 1.5mL tube. These steps were repeated 4 times. The collected supernatant was spun at top speed for 20 min at 4°C and subsequently, protein concentration was measured by BCA assay. 100µg of proteins in 50µL lysate were first incubated with TCEP 5mM for 30 min at 37°C (500rpm) and then with iodoacetamide 10mM in the same conditions and in the dark. 50µL ammonium bicarbonate 100mM was added to dilute urea to 4M before starting proteolysis by LysC (Wako, 0.4µg) at 37°C for 3h at 500rpm. Additional 100µL ammonium bicarbonate 100mM was then added to further dilute urea to 2M. Trypsin (Promega, 1.2µg) was added for proteolysis at 37°C overnight. The reaction was quenched with formic acid 5% and the C18 clean-up (C18 17-170µg, The Nest Group) was performed as described above.

### Mass spectrometry data acquisition

LC-MS/MS analysis was performed on an Orbitrap Exploris480 mass spectrometer (Thermo Fisher) coupled to an Vanquish Neo liquid chromatography system (Thermo Fisher). Peptides were separated using a reverse phase column (75 µm ID x 400 mm New Objective, in-house packed with ReproSil Gold 120 C18, 1.9 µm, Dr. Maisch GmbH) across a linear gradient from 7 to 35% in 120 min (buffer A: 0.1% (v/v) formic acid; buffer B: 0.1% (v/v) formic acid, 80% (v/v) acetonitrile). The DDA data acquisition mode was set to perform one MS1 scan followed by a maximum of 20 dependent scans with MS1 scans (R=60’000 at 400 m/z, normalized AGC (%)=100 and max. IT set to auto), HCD fragmentation (NCE=28%), isolation windows (1.4 m/z) and MS2 scans (R=15’000 at 400 m/z, normalized AGC (%)=200 and max. IT set to auto). A dynamic exclusion of 25s was applied and charge states lower than two and higher than seven were rejected for the isolation.

### Mass spectrometry data analysis

Acquired spectra were searched using the MaxQuant software package version 1.5.2.8 embedded with the Andromeda search engine (99) against the *Saccharomyces cerevisiae* reference dataset (http://www.uniprot.org/, downloaded on 17.05.2021, 6’057 proteins) extended with reverse decoy sequences. The search parameters were set to include a double multiplicity with heavy Lys (Lys8) and a maximum of three labeled amino acids. Semi-tryptic (free N terminal) with a maximum of two missed cleavage were considered. Carbamidomethyl was set as static peptide modification, while oxidation (M) and deamidation (N-ter) as variable modification. The “match between runs” option was set up for the search, and the MS and MS/MS mass tolerance set to 10 ppm. A false discovery rate of < 1% was used for protein identification. The experiment was performed with four independent replicates and only proteins identified in the light and heavy form in at least 2 replicates per condition were considered for the analysis. Protein labeling was then computed as log2(H/(H+L)) where H and L are respectively the intensities of new synthetized proteins and protein intensity before the pulsed. Statistical analysis was performed using unpaired two-sided t-test. The entire dataset, including raw data, generated tables and scripts used for the data analysis are available in the PRIDE repository (PXD046820).

### Tandem Affinity Purification (TAP)

Tandem Affinity Purification (TAP) of pre-ribosomal particles and untagged control cells was performed as described in (101). Briefly, 2 liters of OD_600_ = 3 were harvested and lysed with glass beads and lysis buffer (50mM Tris-HCl pH 7.5, 75mM NaCl, 1.5mM MgCl_2_, 0.15% NP40) in a Planetary mill PULVERISETTE. Lysates were briefly clarified by centrifugation at 4°C to remove debris and further cleared at 18’000 rpm using a SS-34 rotor at 4°C for 15 min. Lysates were incubated with IgG Sepharose beads, before eluting the TAP particles by TEV protease. Eluates were incubated with calmodulin CaM-Sepharose beads, and the TAP particles eluted with elution buffer (10mM Tris-HCl pH 8, 50mM NaCl, 5mM EGTA). Proteins were then precipitated with TCA, resuspended in 1x LDS (Invitrogen, #NP0007), separated on NuPAGE 4-12% Bis-Tris gradient gels (Invitrogen, #NP0321) and analyzed by silver staining or Western blotting.

### In vivo pulse chase biotinylation assay

Cells were grown overnight in 20mL SD complete media without biotin (0.17% (w/v) yeast nitrogen base without amino acids and biotin (Sunrise Science Products, #1523-100), 2% (v/v) glucose, 0.5% (w/v) NH_4_ sulfate and amino acids mix) supplemented with low biotin (5nM). Cells were then diluted to OD_600_= 0.3, and estradiol was added to a final concentration of 5nM for 2 hours. Subsequently, cells were filter washed twice on a 0.45µM pore size nitrocellulose membrane, with an appropriate amount of SD complete low biotin to remove excess estradiol. Cells were resuspended in a flask containing fresh SD media containing 1mM auxin (IAA) and high biotin (10nM) to induce BirA degradation and biotinylation of AVI-tagged proteins. After 1 hour of biotin pulse and BirA degradation, cells were harvested for immunoblotting or immunoprecipitation with Streptavidin magnetic beads (Invitrogen, #65605D). Filter washed cells were transferred to fresh SD complete media with low biotin and samples were collected immediately or after 1 hour (chase step) and further processed for pull-down with Streptavidin conjugated magnetic beads.

### Puromycin incorporation assay

The assay was performed as in (24). Briefly, exponentially growing cells were incubated with puromycin 100µg/mL for 15 minutes at 30°C. TCA was added to stop the reaction at 10% (v/v) final concentration and samples were incubated for 20 minutes on ice. Cells were washed twice with ice-cold acetone and resuspended in 0.2M NaOH PMSF 0.1mM for 10 min on ice. Then 2X Laemmli buffer (100mM Tris-HCl pH 6.8, 20% (v/v) glycerol, 4% (w/v) SDS, fresh 200mM DTT, 0.2% (w/v) Bromophenol Blue) was added and samples were boiled for 15 min before Western blotting analysis.

### Plunge-freezing

Yeast cell cultures were grown overnight in YPD medium on a shaker (180 rpm) at 30°C. Cultures were diluted to OD_600_ = ∼0.05 in fresh YPD medium, grown until OD_600_ = ∼0.3 and then divided. One half was treated with 1 µM of rapamycin (LC laboratories, dissolved in 90% EtOH and 10% Tween) to induce starvation, and the other half with the same volume of EtOH as a negative control (25). Samples were harvested at OD_600_ = ∼0.6 by spinning down for 2 min at 800 *g* at room temperature and adjusted to OD_600_ = ∼3 with conditioned media. 4 µL of cell suspension were then applied two times onto R 2/2 Cu 200 mesh EM grids (specially treated, Quantifoil), which were negatively glow discharged twice (25mA for 45sec). Cells were immediately vitrified by plunge freezing after blotting for 5-6 seconds from the back (Teflon sheet on front pad) into liquid ethane/propane mixture via Vitrobot Mark IV (Thermo Fisher Scientific) in a chamber conditioned to 25 °C and 95% humidity (102).

### Cryo-FIB milling and CryoET analysis

Vitrified cells on EM grids were imaged and subjected to automated sequential FIB milling using a Focused Ion Beam – Scanning Electron Microscope (FIB-SEM; Crossbeam 550, Zeiss) as described (103). Grids were loaded onto a pre-tilted Autogrid holder using a VCM loading station (Leica Microsystems) and transferred using a VCT500 shuttle (Leica Microsystems) to an ACE500 (Leica Microsystems) for tungsten sputter coating (4nm thick) under cryogenic conditions (103, 104). The grids were then transferred using the VCT500 to the FIB-SEM instrument, where an organometallic platinum precursor was applied onto each grid via a gas injection system (GIS). Subsequently, automated FIB-milling was set up targeting for a lamella thickness of 250 nm by sequentially applying currents of 700 pA, 300 pA, and 100 pA for rough milling, and 50 pA for polishing.

The cryo-FIB milled samples were imaged in a Titan Krios 3 (Thermo Fischer Scientific) at an acceleration voltage of 300 kV and equipped with a BioContinuum imaging filter (slit width used: 10 eV) (Gatan) and a K3 direct electron detector (Gatan). Tilt series were collected at 33,000x magnification (pixel size of 2.68 Å) with a defocus value ranging from -3 µm to -8 µm in SerialEM (105) using PACE-TOMO (106). The tilt series were collected in a dose symmetric scheme, with an angular range between -51° and +69° and 3° increment. The cumulative dose for a tilt series for samples was between 141 and 150 e^-^/Å^2^. Frames were aligned using ‘alignframes*’* and tomograms were manually reconstructed in IMOD (107) at a binning factor of 4. The contrast of the reconstructed tomograms (pixel size of 10.71 Å) was further improved by filtering via IsoNet package applying *defocus 5; snrfalloff 1.2; deconvolution strength 1* (87).

### Template matching and concentration estimation for ribosomes

Template matching was performed using Dynamo in MATLAB (108) on contrast tranfer function (CTF) deconvolved tomograms obtained via IsoNet (87). Firstly, ∼2,000 particles from 20 tomograms of the unstressed yeast WT cells were manually picked and sub-tomogram averaging was performed at a binning factor of 4. The averaged ribosome structure was then used as a reference in template matching for the reconstructed tomograms from different conditions. The general workflow of template matching was followed based on the details in dynamo wiki webpage (https://wiki.dynamo.biozentrum.unibas.ch/w/index.php/Walkthrough_for_template_matching), where the threshold (‘mcc’) value applied was 0.4. For each condition, ten tomograms with similar thickness (∼220 nm) were selected for analysis. To estimate the ribosome concentration in tomograms, a custom python script (**Supplementary code 1**) was written to generate serials of boxes with a defined volume and randomly placed inside the cellular cytoplasm. The centers of each box were manually checked to avoid that they overlap with organelles, other cellular structures or were placed outside the cell (including the plasma membrane). For each tomogram, the script performed five independent measurements reflecting how many template matching detected ribosomes were contained in a box of defined volume of 200 nm × 200 nm × 200 nm. Then the concentration of ribosomes per tomogram was calculated by averaging the five measurements and applying the following formula:

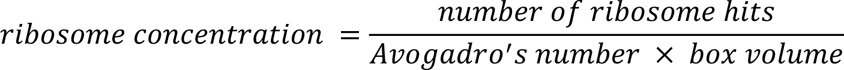

### Live cell microscopy, BiFC measurements and image analysis

Cells were grown overnight in SD media, diluted into fresh growth medium and incubated for ∼ 5h at 30°C to resume exponential phase. Cells were then diluted to OD_600 =_ 0.05 and spotted on a glass bottom plate (Starstedt) functionalized with ConcanavalinA (Sigma, #11028-71-0). Live-cell microscopy was performed using a Nikon Eclipse Ti-E microscope equipped with a Hamamatsu camera, a motorized XY stage and a piezo Z drive and using Micromanager v.1.4.23 software. The incubation chamber of the device was set at 30°C and images were acquired with a 100X oil objectives (NA = 1.45 100x Apo), under non-saturating conditions.

For BiFC assays, a spinning disk confocal fluorescence microscope (Visitron system) equipped with an inverted microscope (Nikon Eclipse TiE), an EM-CCD camera (Andor iXon Ultra, Andor), a motorized XY stage, and a piezo Z drive was used. The system is controlled by VisiVIEW software 5.0.0 (Visitron system). Cells were imaged at 30°C in a temperature-controlled incubator with a ×100 objective lens (NA = 1.4, Nikon CFI Plan Apo).

For quantification, images were processed by Fiji (v. 2.14.0/1.54f) using custom made macros for maximum intensity projections and composites. BiFC quantification of YFP positive cells were carried out using the Fiji plug-in Cell Counter.

### RNA extraction and GelRed staining

Total RNA was extracted with the hot acid phenol method (109). Briefly, 5-10 OD of cells were harvested by centrifugation, washed in RNAse-free water and resuspended in 200µL AE buffer (50mM NaAc, 10mM EDTA), 20µL 10% SDS and 250µL of acid saturated phenol (pH 4.5-5, Roti #A980.1). RNA was extracted at 65°C with shaking, precipitated with ethanol and resuspended in RNAse-free water. 5µg of RNA were mixed with 2.5X sample denaturation mix (65% formamide, 22% formaldehyde, 13% 10X E-buffer), denaturated at 55°C for 15°C, mixed with loading dye (50% Glycerol, 0.3% Xylene cyanol, 0.3% bromophenol blue, 1mM EDTA) and loaded on a 1.2% agarose-6%formaldeyde gel in 1X E-buffer (20mM MOPS, 5mM NaAc, 0.5mM EDTA pH 7.0). RNAs were visualized with Gel Red (Botium #41003) and photographed with a Quantum imager (Vilber). rRNA bands were quantified using Fiji (v. 2.14.0/1.54f) software.

### Protein purification and fluorescence polarization assays

Transformed Rosetta2(DE3) pLysS cells were grown at 37°C till OD_600_= 2.5 in TB medium, cold-shocked for 20 min in ice-water and induced with 0.5 mM IPTG for 16 hours at 30°C. Cells were harvested by centrifugation, resuspended in lysis buffer (GST-buffer containing DNAseI, PMSF, Pepstatin A, Leupeptin and Lysozyme) and lysed by high-pressure homogenization (Emulsiflex). Cleared supernatant was loaded onto a GSTprep 16/60, washed with GST-buffer (25 mM MOPS, pH 7.6; 150 mM NaCl; 1 mM DTT; 5 % (v/v) GlyOH) containing 1 M NaCl and eluted with 10 mM GSH in GST-buffer. GST-tagged Ifh1-AD was fractioned by gel filtration (S75 16/600 pg), an aliquot analyzed by SDS-PAGE, and pure proteins pooled and flash-frozen in liquid nitrogen.

Fluorescence polarization assays were performed in 20 mM MOPS, pH 7.6; 100 mM NaCl; 25 mM D-Trehalose; 1 mM DTT; 1 % (v/v) GlyOH; 0.01 % (v/v) Triton X-100; 0.1 g/ L BSA. Briefly, 25 µL of 40 nM TAMRA-labeled Stm1 IDR-peptide was mixed 1:1 with increasing amounts of GST-Ifh1-AD and incubated for 20 min at room temperature. Fluorescence polarization was measured using a CLARIOStar micro-plate reader (BMG Labtech). The target value of free fluorescent peptide was set to 20 mP. Triplicates were measured with different protein preparations of GST-Ifh1-AD and baseline corrected against a GST-only control. To obtain K_d_ value(s) baseline corrected data were then fit to a one site specific binding model in GraphPad Prism (v10.2.1).

